# The consequences of differential origin licensing dynamics in distinct chromatin environments

**DOI:** 10.1101/2021.06.28.450210

**Authors:** Liu Mei, Katarzyna M. Kedziora, Eun-Ah Song, Jeremy E. Purvis, Jeanette Gowen Cook

**Affiliations:** Department of Biochemistry & Biophysics, University of North Carolina at Chapel Hill, NC 27599; Department of Genetics, University of North Carolina at Chapel Hill, NC 27599; Bioinformatics and Analytics Research Collaborative (BARC), University of North Carolina at Chapel Hill, NC 27599

## Abstract

MCM complexes are loaded onto chromosomes to license DNA replication origins in G1 phase of the cell cycle, but it is not yet known how mammalian MCM complexes are adequately distributed to both euchromatin and heterochromatin. To address this question, we combined time-lapse live-cell imaging with fixed cell immunofluorescence imaging of single human cells to quantify the relative rates of MCM loading in heterochromatin and euchromatin at different times within G1. We report here that MCM loading in euchromatin is faster than in heterochromatin in very early G1, but surprisingly, heterochromatin loading accelerates relative to euchromatin loading in middle and late G1. These different loading dynamics require ORCA-dependent differences in ORC distribution during G1. A consequence of heterochromatin origin licensing dynamics is that cells experiencing a truncated G1 phase from premature cyclin E expression enter S phase with under-licensed heterochromatin, and DNA damage accumulates preferentially in heterochromatin in the subsequent S/G2 phase. Thus G1 length is critical for sufficient MCM loading, particularly in heterochromatin, to ensure complete genome duplication and to maintain genome stability.

## INTRODUCTION

Eukaryotic cell cycle progression is a highly orchestrated and strictly-regulated process. One key event during the cell cycle is DNA replication, and it must be tightly controlled to ensure complete and precise genome duplication (for reviews, see ^1, 2^). DNA replication in mammalian cells initiates at discrete sites called replication origins that are not strictly defined by DNA sequence, but rather, by other aspects of chromatin ^3–5^. In G1 phase, origin DNA is “licensed” by the loading of MCM complexes that will later be activated in S phase to form the core of the replicative helicase ^6–8^. Successful genome duplication requires many active DNA replication origins per chromosome; in mammalian cells thousands of origins are licensed in each G1, and then a subset are activated, or “fired” in S phase ^9^. Origins that are licensed but not fired are dormant origins, and they are induced to fire near stalled forks in order to ensure complete replication ^10, 11^. When too few licensed origins are available in a local genomic region, the resulting incomplete replication promotes chromosome breaks and genome instability ^12, 13^. Thus, successful replication requires sufficient origin licensing in all genomic regions.

Chromosomes are not uniform substrates for replication however, and chromatin structure and DNA accessibility vary widely among different genomic regions. If MCM loading is too unevenly distributed, then some regions become vulnerable to under-replication. For example, common fragile sites are more likely to be under-replicated in part because of large inter-origin distances that flank these sites and few available licensed dormant origins ^14–16^. The reasons for large distances between origins near common fragile sites include sparse MCM loading during the preceding G1^16^, but what causes regions of low MCM loading are not yet fully understood. One possibility is that some regions are licensed slowly or later during G1, and these differential licensing dynamics create a risk for local under-licensing when S phase begins. To date, little is known about MCM loading dynamics during G1 in any system. Are all genomic regions licensed at the same speed, or is there a temporal hierarchy among regions? If licensing proceeds by a preferred temporal order, then are regions that are licensed last more likely to be under-licensed and then under-replicated? We hypothesized that different chromatin environments influence licensing dynamics, and that those dynamics, in turn, impact DNA replication.

Chromatin can be divided into two distinct environments, heterochromatin and euchromatin. Euchromatin is loosely packed and associated with transcriptional activity, whereas heterochromatin is more condensed and generally transcriptionally repressed ^17–19^. Thus far, the link between overall MCM loading and chromatin states has primarily been explored in the context of replication timing studies. Origins fire at different times in S phase, and a general correlation between less accessible heterochromatin and replication later in S phase has been reported in many species and cell types ^20–22^. Replication timing programs have been suggested as mechanisms that avoid replication stress and maintain genome and chromatin organization ^23–25^, but the relationship between G1 phase dynamics and S phase dynamics are still unknown. The relative local amounts of loaded MCM complexes and MCM loading factors, such as the Origin Recognition Complex, have been implicated in establishing replication timing programs within S phase ^26^. Although replication timing correlates with this MCM loading density, timing within S phase is not a direct measure of origin licensing because origin firing time can be influenced by other factors in S phase ^20, 27, 28^, nor does timing analysis itself explain how differences in MCM loading arise during G1 phase.

It is also currently unknown how MCM loading is adequately distributed during G1 phase to ensure full genome duplication. Since heterochromatin is considered a repressive and less accessible environment, MCM loading may be less efficient in heterochromatin compared to loading in euchromatin. MCM and its loading factor ORC are indeed more concentrated in genomic regions associated with active chromatin marks, suggesting that euchromatin is particularly permissive for origin licensing, although the precise mechanisms driving licensing enrichment in euchromatin are not yet clear ^29^. Nonetheless, some mechanistic links between MCM loading efficiency and different chromatin features or chromatin modifying enzymes are known ^16, 30–32^. MCM loading is generally enriched at sites with low nucleosome density ^16, 33, 34^. Several chromatin-modifying factors are reported to generally promote MCM loading, such as the chromatin remodeler SNF2H ^35^, the HBO1 histone acetylase ^36, 37^, the histone variant H2A.Z ^38^, the histone H4 lysine20 methyltransferase PR-Set7 ^39^, and the heterochromatin binding protein ORCA ^40, 41^. In contrast, the Sir2 histone deacetylase suppresses MCM loading at some budding yeast origins ^42^. However, these studies did not address the dynamics or distribution of mammalian MCM loading during G1.

Although euchromatin is efficiently licensed, a substantial fraction of mammalian DNA resides in heterochromatin that must also be replicated each cell cycle. In mammalian cells, the longest inter-origin lengths typically observed were ~400-600 kb ^43, 44^, yet heterochromatin regions can be as long as multiple megabases, including the extreme example of the inactive X chromosome which is largely heterochromatic, but is still replicated every S phase ^45, 46^. Thus, at least some origins in heterochromatin must still be licensed during G1 to ensure complete replication in S phase. Importantly, once S phase begins, the MCM loading factors (ORC, CDC6, and CDT1) are degraded or inactivated to prevent any more MCM loading after the end of G1 ^47–49^. This strict separation of origin licensing and origin firing avoids re-replication, a source of endogenous DNA damage and genome instability ^50–52^. An important consequence of blocking MCM loading factors in S phase is that all of the MCM loading needed for a complete S phase must occur before the G1/S transition. Thus, the amount and distribution of MCM loading at the end of G1 phase determines the likelihood of a successful S phase ^53^. However, it is still unknown how the entire genome – both euchromatin *<u>and</u>* heterochromatin - receives sufficient MCM before S phase starts.

Here, we combined live cell imaging with fixed cell imaging to quantify MCM loading in both heterochromatin and euchromatin as G1 progresses. We discovered that the MCM loading rate is higher in euchromatin than in heterochromatin during early G1. Loading in both chromatin types accelerates during G1 phase, but heterochromatin loading accelerates *more* than euchromatin does to achieve similar rates and concentrations by the end of G1. Because MCM loading in heterochromatin is later during G1, and heterochromatin typically replicates later in S phase, cells that start S phase prematurely experience under-replication and DNA damage preferentially in heterochromatin. These findings quantify MCM loading dynamics with high temporal resolution to reveal a source of unique vulnerability to genome instability specifically in heterochromatin.

## MATERIALS AND METHODS

### Cell culture

HEK293T and RPE1-hTERT cells were originally obtained from the ATCC and confirmed to be mycoplasma negative. HEK293T, and RPE1-hTERT were cultured in Dulbecco’s Modified Eagle Medium (DMEM) supplemented with 2 mM L-glutamine and 10% fetal bovine serum (FBS) and incubated in 5% CO2 at 37°C. All cell lines were authenticated by STR profiling (Genetica Cell Line Service, Burlington, NC), were passaged with trypsin, and not allowed to reach confluence.

### Cloning

All constructs were generated using either the Gateway cloning method or by Gibson Assembly following standard protocols as described before ^54^. PCR fragments were amplified using Q5 polymerase (New England Biolabs, NEB). DNA fragments were isolated using the Qiaprep spin miniprep kit (Qiagen). Plasmids were transformed into either DH5α or Stbl2 *Escherichia coli* strains for propagation. pENTR constructs were combined with the expression constructs: pInducer 20 (Addgene plasmids#44012). Plasmids were validated via sequencing (Eton Biosciences) for the desired insert using appropriate primers. YFP tagged ORC1 is a gift from Supriya Prasanth ^55^, and was cloned to pInducer20-neo using Gateway cloning. The CDK activity reporter plasmid CSII-EF zeo DHB-mCherry was a gift from S. Spencer (University of Colorado-Boulder, Boulder, CO).

### Cell line construction and inducible protein production

To package lentivirus, pInducer20-mVenus-MCM3, pInducer20-YFP-ORC1, or pInducer20-Cyclin E1 ^54^, were co-transfected with ΔNRF and VSVG plasmids (gift from Dr. J. Bear) into HEK293T using 50 μg/mL Polyethylenimine-Max (Aldrich Chemistry). Viral supernatants were transduced with 8 ug/mL polybrene (Millipore, Burlington, MA) into RPE1-hTERT cells for 24 hr. Transduced cells were selected with 500 ug/mL neomycin (Gibco) for 1 week. 500 cells were seeded in 15 cm dish and individual positive clones were hand-picked and screened by immunoblotting and evaluated by flow cytometry. To overproduce Cyclin E1, cells were treated with 100 ng/mL doxycycline (CalBiochem, San Diego, CA) for 7 hr or 18 hr in complete medium. To overproduce mVenus-MCM3, cells were treated with 500 ng/mL doxycycline for 2 hr in 10% FBS, DMEM, L-glutamine. To overproduce YFP-ORC1, cells were treated with 100 ng/mL doxycycline for 24 hr in complete medium. Control cells were in complete medium without doxycycline.

### Live cell imaging

Cells were plated on fibronectin (1 ug/cm^2^, Sigma) coated F1141 glass-bottom plates (Cellvis) with FluoroBrite™ DMEM (Invitrogen) supplemented with 10% FBS, 4 mM l-glutamine, and penicillin/streptomycin. Fluorescence images were acquired using a Nikon Ti Eclipse inverted microscope with Plan Apochromat dry objective lenses 20x (NA 0.75) or 40x (NA 0.95). Images were captured using a Andor Zyla 4.2 sCMOS detector with 12 bit resolution. Autofocus was provided by the Nikon Perfect Focus System (PFS), and a custom enclosure (Okolabs) was used to maintain constant temperature (37°C) and atmosphere (5% CO2) in a humidified chamber. All filter sets were from Chroma, CFP – 436/ 20 nm; 455 nm; 480/40 nm (excitation; beam splitter; emission filter), YFP – 500/20 nm; 515 nm; 535/ 30 nm; and mCherry – 560/40 nm; 585 nm; 630/ 75 nm. Images were collected every 10 minutes using NIS-Elements AR software. No photobleaching or phototoxicity was observed in cells imaged by this protocol.

### Tracking and segmentation

Individual cells were segmented and tracked in time-lapse movies by a user-assisted approach as previous described ^56^. In brief, all movies were pre-processed using rolling ball background subtraction. Individual cells in the movie were manually tracked using a set of in-house developed ImageJ scripts. Using user-defined tracks, nuclear regions of interest (ROI) were segmented automatically based on intensity of PCNA followed by separation of touching nuclei by a watershed algorithm. In case of failed segmentation, the user could manually define polygons as replacement ROIs. The same set of ROIs was used to analyze all fluorescence channels.

### PCNA variance and CDK activity

PCNA variance was quantified as described previously ^56^. The PCNA pattern was analyzed within nuclear ROIs, then images were processed in a series of steps implemented in Fiji (ImageJ): 1. Images procession: Image smoothing, edge enhancement and nuclear regions reduction. 2. Quantification of processed PCNA signal, a sum of mean and standard deviation of variance image showed the highest contrast at the beginning and the end of the S phase and was therefore used for cell cycle phase delineation. CDK1/2 activity was quantified as the ratio of cytoplasmic to nuclear mean intensity of the DHB-mCherry sensor (cytoplasm quantified in a 15-pixel ring outside the nuclear segmentation).

### siRNA transfections

For siRNA treatment of RPE cells, Dharmafect 4 (Dharmacon) was mixed in Optimem (Gibco) with the appropriate siRNA according to manufacturer’s instructions, then diluted with DMEM, 10% FBS, and L-glutamine and added to cells after aspirating old media. The next day, the siRNA mix was aspirated and replaced with fresh DMEM, 10% FBS, L-glutamine, collecting samples 48 hr after the start of siRNA treatment. Generally, the siRNA were siControl (Luciferase) at 100 nM or a mixture of two MCM3 siRNA (2859 and 2936 at 100 nM each) or siORCA (100 nM). The luciferase siRNA and MCM siRNA were synthesized by Sigma ^54^. The ORCA siRNA was synthesized by Dharmacon ^41^.

~~~
siControl (Luciferase)- cuuacgcugaguacuucga
siMCM3-2859 5’- augacuauugcaucuucauug
siMCM3-2936 5’- aacauaugacuucugaguacu
siORCA 5’- ccaaccaggacuacgaauu
~~~

### Immunofluorescence for chromatin associated proteins

Cells were washed with phosphate-buffered saline (PBS) immediately after live cell imaging, CSK buffer (300 mM sucrose, 300 mM NaCl, 3 mM MgCl^2^, 10 mM PIPES pH 7.0) with 0.5% triton x-100 and protease and phosphatase inhibitors (0.1 mM AEBSF, 1 µg/ mL pepstatin A, 1 µg/ mL leupeptin, 1 µg/ mL aprotinin, 10 µg/ ml phosvitin, 1 mM β-glycerol phosphate, 1 mM Na-orthovanadate) was added to each well for 5 min on ice. Then CSK buffer with soluble proteins was discarded, and cells were fixed with 1 ml 4% paraformaldehyde for 15 min and washed with PBS twice. Cells were blocked in 1% bovine serum albumin in PBS for 1 hr, and incubated with the primary antibody overnight at 4°C. Primary antibodies used were: Mcm2 (1:1000, BD Biosciences, Cat#610700), Mcm3 (1:1000, Bethyl Laboratories, Cat#A300-192A), histone H3K9me3 (1:5000, Active Motif, Cat#39062), histone H3K9me3 (1:3000, Active Motif, Cat#61014), HP1 (1:500, Santa Cruz Biotechnology, Cat#sc-515341), histone H4ac (1:3000, Active Motif, Cat#39244), ORC4 (1:100, Santa Cruz Biotechnology, CA, Cat#136331), GFP (1:1000, Invitrogen, Cat#A11122), 53BP1 (1:1000, Novus, Cat#NB100304), RPA (1:200, Cell Signaling Technology, Cat#2208). Cells were incubated with secondary Alexa Fluor series antibodies (all 1:500, Invitrogen) for 1 hr at room temperature and then with 1 µg/ml DAPI for 5 min. Secondary antibodies used were: donkey anti-mouse-Alexa 488 (Jackson ImmunoResearch), donkey anti-rabbit-Alexa 594 (Jackson ImmunoResearch), donkey anti-rabbit-Alexa 488 (Jackson ImmunoResearch), goat anti-mouse-Alexa 594 (Jackson ImmunoResearch), Donkey anti-rat-Alexa 647 (Jackson ImmunoResearch). Z stack images were collected using a Zeiss 880 upright confocal microscope with a 63 oil-immersion objective lens (Pln Apo 63x/1.4 numerical aperture [NA]); The pixel size is 0.07µM and the xy resolution is 380 × 380 for each slice. the distance between two slices is 0.2 µM. Images were acquired in an automated fashion with the ZEN acquisition software (Ceiss). No photobleaching was observed during acquisition of the stacks.

### Single cell analysis of 3D confocal images

Otsu thresholding was performed on each full nucleus using the DAPI channel. Image segmentation was performed using an Image J script. Heterochromatin was thresholded to the 10%, 20% or 50% brightest HP1 or H3K9me3 pixels in the nuclear mask. All fluorescence signals for each channel were calculated using custom Python scripts (v3.7.1) in Jupyter Notebooks (v6.1.4). RPA foci were counted within nuclear ROIs in the channel to be quantified using the 3D Objects Counter plugin in ImageJ. Data were visualized using Jupyter Notebooks (Python graphical libraries Matplotlib ^57^ and Seaborn ^58^) and GraphPad Prism (v8).

### Total lysate and chromatin fractionation

Cells were collected by trypsinization. Total protein lysates for immunoblotting were generated as described previously ^54^, cells were lysed on ice for 20 min in CSK buffer with 0.5% triton x-100 and protease and phosphatase inhibitors. Cell lysates were centrifuged, and the supernatants kept for a total protein Bradford Assay (Biorad, Hercules, CA) using a BSA standard curve. Chromatin fractionation for immunoblotting was performed as described previously ^54^ using CSK buffer with 1 mM ATP, 5 mM CaCl_2_, 0.5% triton x-100, and protease and phosphatase inhibitors to isolate insoluble proteins and S7 nuclease (Roche) to release DNA bound proteins. A protein assay was performed for chromatin fraction quantification.

### Immunoblotting

Samples were diluted with SDS loading buffer and boiled. Proteins were separated on SDS-polyacrylamide-gels, then transferred onto polyvinylidene difluoride (PVDF) membranes (Thermo Fisher, Waltham, MA). Membranes were blocked in 5% milk in Tris-Buffered-Saline-0.1%-tween-20 (TBST) at room temperature for 1 hr. Then membranes were incubated in primary antibody overnight at 4°C in 2.5 % milk in TBST. The next day, blots were washed with TBST then incubated in secondary antibody conjugated to horseradish peroxidase in 2.5% milk in TBST for 1 hr, washed with TBST for 3 times. For detection, membranes were incubated with ECL Prime (Amersham, United Kingdom) and exposed to autoradiography film (Denville, Holliston, MA) or detected by a ChemiDoc imaging system (Biorad). Ponceau S staining for total protein (Sigma Aldrich) was typically used as loading control. The following primary antibodies were used for immunoblotting: Mcm2 (BD Biosciences, San Jose, CA, Cat#610700), Mcm3 (Bethyl Laboratories, Montgomery, TX, Cat#A300-192A), Cdt1 (Santa Cruz, CA, Cat#sc-365305), histone H3 (Gene Script, NJ, Cat#A01502), histone H3ac (MilliporeSigma, CA, Cat#06-599), histone H3K9me3 (Active motif, CA, Cat#39062), α-tubulin (Sigma Aldrich, CA, Cat#9026). PCNA (Santa Cruz Biotechnology, Cat#sc-25280), HP1 (Santa Cruz Biotechnology, Cat#sc-515341). The secondary antibodies were used for immunoblotting: anti-rabbit IgG HRP-conjugated (1:10000, Jackson Immuno Research), goat anti-mouse IgG HRP-conjugated (1:10000, Jackson Immuno Research).

### Flow cytometry

Analysis of chromatin bound MCM was performed as described in Matson *et al*. ^54^. Briefly, cells were treated with 10 μM EdU (Santa Cruz Biotechnology) for 30 min before collection. Cells were collected then lysed on ice for 8 min in CSK buffer (10 mM Pipes pH 7.0, 300 mM sucrose, 100 mM NaCl, 3 mM MgCl_2_) with 0.5% triton x-100 with protease and phosphatase inhibitors. Cells were washed with 1% BSA-PBS then fixed in 4% paraformaldehyde in PBS for 15 min at room temperature. For EdU detection, samples were incubated in PBS with 1 mM CuSO_4_, 1 μM Alexa 647-azide (Life Technologies), and 100 mM ascorbic acid (fresh) for 30 min at room temperature in the dark. Then cells were labeled in primary antibody and incubated at 37°C for 1 hr in the dark. Next, cells were resuspended in secondary antibody and incubated at 37°C for 1 hr in the dark. Finally, cells were resuspended in 1% BSA-PBS +0.1% NP-40 with 1 μg/mL DAPI (Life Technologies) and 100 μg/mL RNAse A (Sigma Aldrich) and incubated overnight at 4°C in the dark. Data were collected on an Attune NxT flow cytometer (Thermo Fisher Scientific) and analyzed using FCS Express 7 (De Novo Software) software. Control samples were prepared omitting primary antibody or EdU detection to define thresholds of detection as in Matson et al 2017 ^54^. The following antibodies were used: primary: Mcm2 (1:100, BD Biosciences, Cat#610700) or ORC4 (1:50, Santa Cruz Biotechnology, Cat#136331), secondary: Donkey anti-mouse-Alexa 488 (Jackson ImmunoResearch),

### Cell synchronization and treatments

To synchronize cells in quiescence (G0), RPE1-hTert cells were grown to 100% confluency and incubated for another 48 hr in 10% FBS, DMEM, L-glutamine. G0 cells were then released by passaging 1:10 with trypsin to new dishes in 10% FBS, DMEM, L-glutamine, cells were collected at different timepoints (16 hr, 18 hr, 20 hr, 23 hr, 26 hr, 28hr, 30 hr) to enrich for different phases of cell cycle. For aphidicolin (Sigma) treatment, asynchronous RPE cells were treated with 25 ng/mL or vehicle control for 4 hr.

### Statistical analysis

Bar graphs represent means, and error bars indicate standard error of the mean (SEM), unless otherwise noted. The number and type of replicates are indicated in the figure legends. Significance tests were performed using a one-way ANOVA test, as indicated in the figure legends, unless otherwise specified. Statistical significance is indicated as asterisks in figures: * P ≤ 0.05, ** P ≤ 0.01, *** P ≤ 0.001 and **** P ≤ 0.0001. GraphPad Prism v.8.0 and python were used for statistical analysis.

## RESULTS

### An experimental system for analyzing subnuclear MCM loading dynamics within G1 phase

MCM complexes can begin to load during telophase of mitosis, and loading continues through G1 phase, peaking just before the G1/S transition; MCM complexes are unloaded during S phase ^59, 60^. The total levels of MCM subunits are constant throughout the cell cycle, but the levels of loaded MCM change as cells progress through the cell cycle ^61^. To quantify and analyze the dynamics of MCM loading in G1 phase, we developed a method to correlate intranuclear MCM loading positions in individual human cells with cell cycle progression (**Figure 1A, B**). We created derivatives of the human retinal pigmented epithelial cell line, RPE1-hTert, that harbor reporters of S phase entry and CDK1/CDK2 activity. We performed time-lapse imaging on these asynchronously proliferating cells to define the cell cycle age of individual cells (time elapsed since the previous mitosis), cell cycle phase (G1, S, G2, M), and CDK1/2 activity for each cell. We extracted the cells with nonionic detergent in the presence of 300 mM salt to remove soluble MCM immediately after the movie was stopped. We then fixed the remaining chromatin-bound proteins for immunofluorescence (IF) using anti-MCM3 antibody as a marker of the MCM2-7 complex, anti-HP1 antibody as a marker of heterochromatin ^62, 63^, and DAPI for DNA content (**Figure 1B**). We performed confocal imaging and quantified the colocalization of loaded MCM and HP1 in G1 cells (**Supplementary Video 1**). This approach is suitable for analyzing unperturbed proliferating populations to identify cell cycle-related changes that would be difficult to detect by immunoblotting or flow cytometry. Non-extractable, salt-resistant MCM complexes are strongly correlated with DNA replication origins and replication competence in vitro ^64, 65^ and in vivo ^54, 66^.

**Figure 1.**
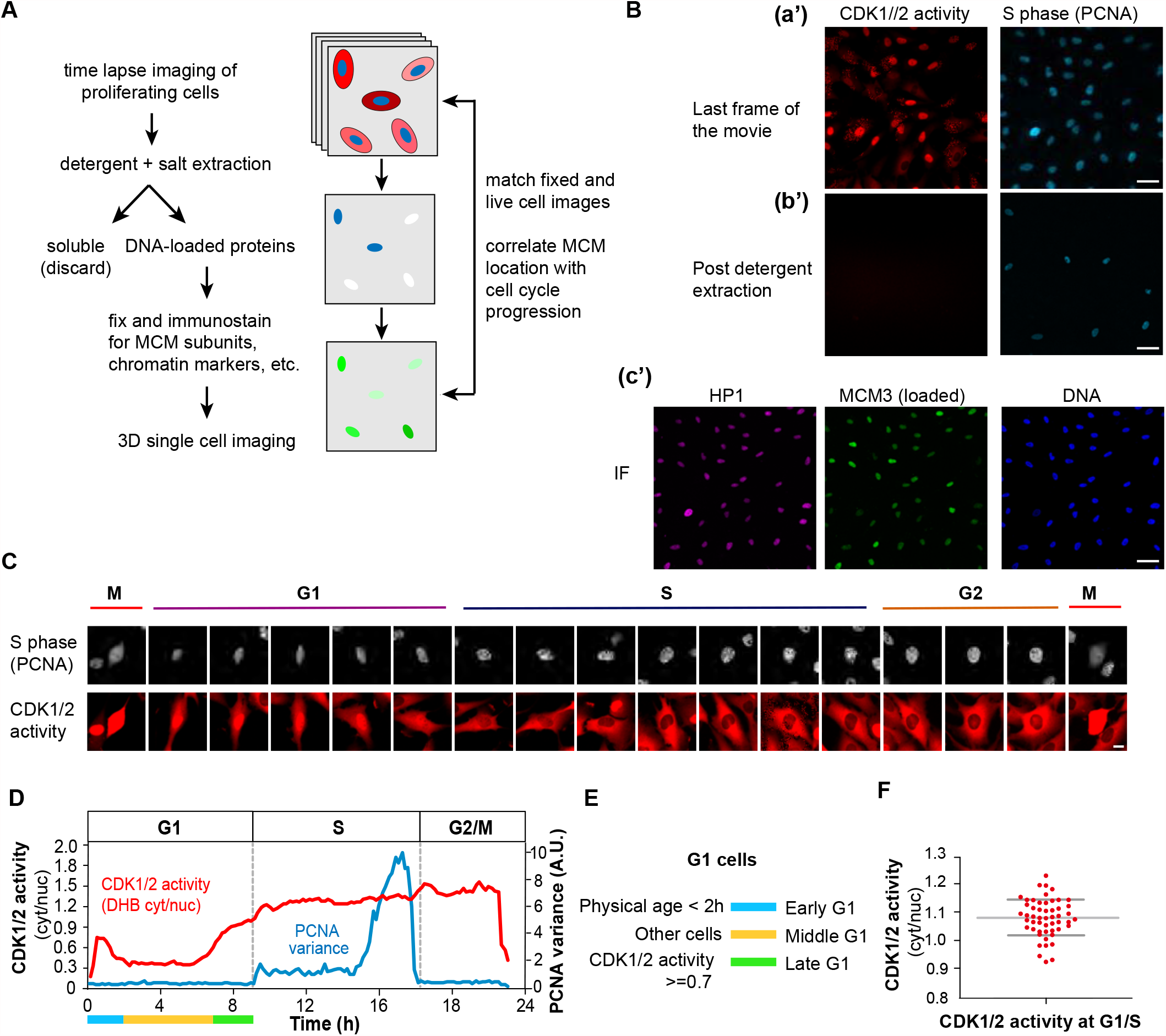
An experimental system for analyzing subnuclear MCM loading dynamics within G1 phase. (**A**) Workflow. RPE-hTert cells were subjected to live cell time-lapse imaging, then soluble proteins were extracted with non-ionic detergent and salt, and cells were fixed immediately after imaging for confocal immunofluorescence staining. (**B**) Representative example of combining live cell imaging with fixed cell imaging. (a’) Last frames from wide field time-lapse imaging of cells expressing CDK1/2 activity and S phase reporters. (b’) Images collected with the same microscope settings after detergent extraction and fixation; scale bar represents 100 μm. (c’) Immunofluorescence of fixed cells after live cell imaging. Cells were stained for bound HP1 (heterochromatin marker) and loaded MCM3 (MCM2-7 complex marker) and imaged by confocal microscopy; scale bar 100 μm. (**C**) Selected images from wide field time-lapse imaging of one cell. Images were captured every 10 min for 1 cell cycle, and selected frames from one of 50 cells are shown. The scale bar is 10 µm and applies to all images. Images were brightness/contrast adjusted. (**D**) An individual cell trace of PCNA variance and CDK1/2 activity for one cell. CDK1/2 activity is the ratio of mean cytoplasmic DHB-mCherry reporter fluorescence divided by mean nuclear DHB-mCherry fluorescence. Hours are time since mitosis. (**E**) Defining G1 subphases by both physical age and CDK1/2 activity. G1 cells younger than 2 hours after mitosis are early G1 cells; G1 cells older than 2 hours with CDK activity less than 0.7 are middle G1 phase; G1 phase cells with CDK activity equal to or more than 0.7 but not yet in S phase by PCNA variance are late G1 cells. (**F**) Quantification of CDK1/2 activity at the G1/S transition as defined by the increase in PCNA variance. *n* = 50, mean with SEM.

As controls, we validated the MCM3 antibody specificity by the loss of immunostaining of both total and loaded MCM3 in MCM3-depleted cells (**Supplementary Figure S1A and S1B**). Anti-MCM3 staining also showed strong colocalization with MCM2 which is expected from two subunits of the MCM2-7 complex (**Supplementary Figure S1C**). MCM3 also co-localized with an ectopically-expressed YFP-ORC1 fusion in G1 cells indicating that MCM detected this way is at sites of origin licensing (**Supplementary Figure S1D**). Furthermore, our HP1 staining was mutually exclusive with a marker of euchromatin, histone H4 acetylation (H4ac) and colocalized with histone H3 lysine 9 trimethylation, another established heterochromatin mark (**Supplementary Figure S1E-G**).

We defined G1 cells in the asynchronously proliferating population by identifying cells post-mitosis and pre-S phase using the variance of PCNA-mTurq2 localization as previously described ^56^. PCNA-mTurq2 was present throughout the cell cycle, diffusely distributed in G1 nuclei, punctate during S phase, and diffuse again in G2 ^67^. To automatically detect S phase boundaries, we calculated the variance of PCNA intensity across the nucleus. The rapid increase in PCNA variance indicates the onset of S phase, whereas the steep drop in PCNA variance indicates the end of S phase (**Figure 1C and 1D**). In normal RPE1-hTert cells, G1 length varies from 5 hours to 12 hours, with most cells spending 7-8 hours in G1 ^68^. We defined cells younger than 12 hours with no significant increase in PCNA variance as G1 cells. As a measure of CDK2 activity, we monitored the relative localization of a model CDK substrate, DHB-mCherry. CDK2 is activated by cyclin E during G1, and CDK-dependent DHB-mCherry reporter phosphorylation induces translocation from the nucleus to the cytoplasm ^69^. Based on the normal timing of CDK2 and CDK1 activation and previous reports, we attribute reporter translocation in late G1 and early S phase to CDK2 activity and G2 phase reporter localization to a combination of CDK2 and CDK1 activity ^70^; the reporter is not responsive to CDK4 or CDK6 activity ^69, 71^. We quantified the ratio of cytoplasmic to nuclear reporter localization throughout the cell cycle (**Figure 1D**). We tracked complete cell cycles of 50 cells and quantified their CDK1/2 activity when they entered S phase; the cytoplasmic:nuclear localization range was from 0.9 to 1.3, and the majority of cells enter S phase with the CDK1/2 activity around 1.1 (**Figure 1D, F**).

To analyze MCM loading dynamics in different G1 subphases, we categorized G1 cells into three groups according to both their physical age in hours and their CDK1/2 activity. Because the nuclear membrane is undergoing maturation in early G1, and many nuclear components (including the CDK reporter) are still partly cytoplasmic, we categorized early G1 cells by their physical age; cells younger than 2 hours are early G1 cells. However, the large intercellular variation in G1 length meant that CDK activity correlated with G1 progression better than physical age in middle and late G1 phases. Of note, CDK activity increases non-linearly over time in G1 (**Figure 1D**). For example, sisters with different G1 lengths can have very different CDK activities and MCM loading amounts at the same physical age **(Supplementary Figure S2A**). We therefore used CDK activity to define the “molecular age” of middle and late G1 cells instead of simply dividing G1 phase according to time since mitosis. Previous studies have measured CDK1/2 reporter ~0.7 near the point in G1 when cells are committed to S phase, so we chose 0.7 as a mark between middle and late G1 ^69^,^70^. G1 cells older than 2 hours but with CDK1/2 activity still below 0.7 are middle G1 phase cells, and G1 phase cells with CDK1/2 activity above 0.7 are late G1 cells (**Figure 1E**).

### Differential dynamics of MCM loading in euchromatin and heterochromatin

By the analysis outlined above, MCM signal was very low on chromatin in G2 cells, but total nuclear MCM loading signal increased throughout G1 (**Figure 2A, B, Supplementary Figure S2B**). We hypothesized that the dynamics of intranuclear MCM loading during G1 phase are affected by chromatin environments. To test this idea, we used HP1 immunostaining to distinguish heterochromatin from euchromatin and quantified loaded MCM in heterochromatin and euchromatin as a function of CDK activity. As expected, the HP1 signal was unevenly distributed in nuclei and was concentrated in discrete foci ^72, 73^ (**Figure 2A)**. It has been reported that for many eukaryotes, heterochromatin constitutes 10–20% of the genome ^74, 75^. These reports prompted us to set an initial threshold for identifying heterochromatin in confocal images as the pixels with the 20% brightest HP1 signals, whereas euchromatin is the remaining 80% of nuclear pixels. The HP1 locations were more intensely stained with DAPI which is an indicator of condensed chromatin (**Supplementary Figure 2E**). Neither total HP1 nor the distribution of HP1 on heterochromatin and euchromatin changed during G1 phase (**Supplementary Figure S2C, D**).

**Figure 2.**
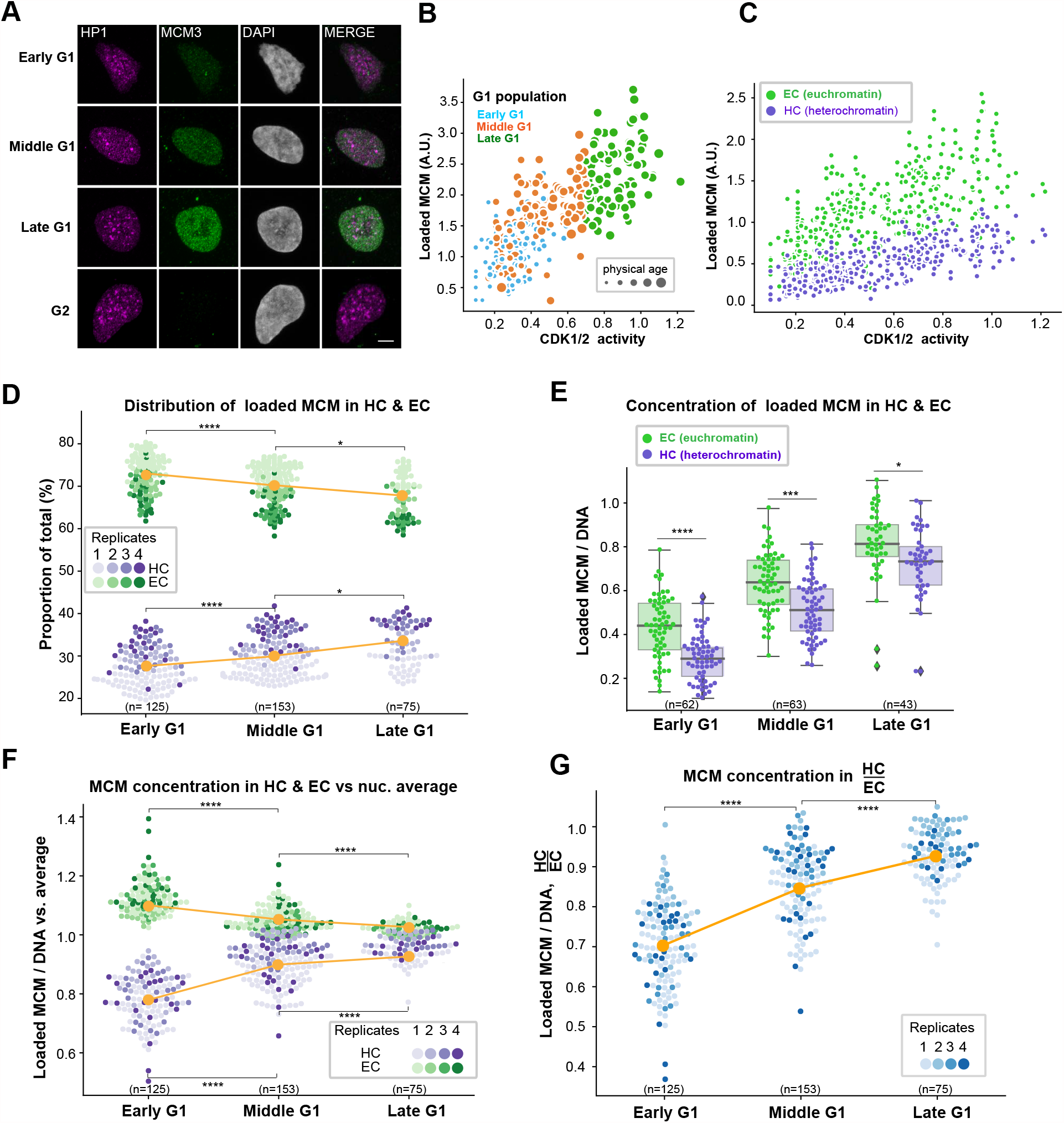
Differential dynamics of MCM loading in euchromatin and heterochromatin. (**A**) Projections of 3D immunofluorescence images of representative cells after live cell imaging as in Figure 1; endogenous HP1 (magenta), endogenous MCM3 (green), DNA stained with DAPI (grey), scale bar 5 μm. (**B**) Loaded MCM3 immunofluorescence signal (y-axis) relative to CDK1/2 activity defined by the cytoplasmic vs. nuclear localization of the reporter (x-axis). Cells are color-coded for early (blue), middle (orange) and late (green) G1 cells defined in Figure 1. The size of data points for single cells correspond to time since mitosis. (**C**) Loaded MCM3 colocalized with HP1 as a marker of heterochromatin (purple dots) and euchromatin (low HP1 regions, green dots) within each cell relative to CDK1/2 activity. All five replicates are shown, n=446. (**D**) Distribution of loaded MCM3 in heterochromatin (purples) and euchromatin (greens) in early, middle and late G1 cells as proportions of the total MCM signal per cell. Four replicates are shown in different shades; means are plotted in orange. One-way ANOVA, Tukey post-hoc test, n (number of cells) is indicated in the figure. (**E**) MCM3 concentration normalized to DNA/DAPI in heterochromatin (purple) and euchromatin (green) in G1 subphases. Box plots show median (solid line) and interquartile ranges (box ends), whiskers mark the minimum or maximum. One-way ANOVA test, five replicates, n (number of cells) is indicated in the figure. (**F**) MCM concentration in heterochromatin or euchromatin relative to the average loaded MCM concentration in whole nuclei; mean is plotted in orange. One-way ANOVA, Tukey post-hoc test, n (number of cells) is indicated in the figure, Four replicates are shown with different shades. (**G**) Ratio of MCM3 concentration in heterochromatin to euchromatin; mean is plotted in orange. One-way ANOVA, Tukey post-hoc test, n (number of cells) is indicated in the figure, Four replicates are shown with different shades. In all panels p value ranges are indicated as * p ≤ 0.05, ** p ≤ 0.01, *** p ≤ 0.001 and **** p ≤ 0.0001.

Our aggregate analysis of cells from multiple independent experiments revealed intercellular variability in total MCM loading signal (the sum of euchromatin and heterochromatin) among cells with similar CDK activity (**Figure 2B)**. Within this heterogeneity, we nonetheless measured distinct and reproducible trends in MCM loading when we analyzed heterochromatin and euchromatin separately (**Figure 2C)**. For both euchromatin and heterochromatin, MCM loading increases throughout G1 indicating a positive loading rate for the entire G1 phase. The total MCM signal is higher in euchromatin than in heterochromatin in keeping with their respective 80%/20% shares of nuclear volume.

Although the MCM loading signal increased for both euchromatin and heterochromatin, the relative *<u>proportion</u>* of MCM loading in heterochromatin vs. euchromatin changed during G1 (**Figure 2D**); chromatin itself did not change during G1 (**Supplementary Figure 2C, D**). We classified G1 subphases by the same criteria defined in **Figure 1D** and **1E**, and found that in early G1, euchromatin accounted for a larger proportion of the total MCM that had been loaded than it did by middle and late G1. As a result, the relative proportions of loading on the two chromatin types converged between early and late G1 so that the difference between them was smaller (**Figure 2D**). This change in proportion plus the general increase in both chromatin types throughout G1 suggested that heterochromatin loading increases faster in middle and late G1 than euchromatin loading.

A feature of heterochromatin is its higher chromatin density, i.e., more DNA and associated protein content per unit volume ^76, 77^. The intensity of DAPI-stained DNA was indeed higher in the regions with the 20% brightest HP1 signals (**Supplementary Figure S2E**). This observation supports our classification of HP1 localization as generally marking heterochromatin, although we acknowledge that HP1 can also be found in euchromatic locations ^63^. To fairly compare MCM loading in heterochromatin to euchromatin, we normalized MCM loading to DNA content at each location to derive the *concentration* of loaded MCM per unit DNA (Loaded MCM / DNA). Similar to measurements of total loaded MCM, the overall concentration of loaded MCM increased during G1 progression (**Figure 2E, Supplementary Figure S2F**). Interestingly, the concentration of loaded MCM on heterochromatin was lower than on euchromatin in early G1, but the difference between them had narrowed by the end of G1 phase (**Figure 2E**). As another means to visualize these differences, we compared the concentration of loaded MCM in heterochromatin and euchromatin to the average concentration of loaded MCM across each whole nucleus. Euchromatin loading starts out higher than the nuclear average early in G1, but the euchromatin and heterochromatin concentrations at their respective locations approach each other by the end of G1 (**Figure 2F**). We observed similar dynamics using another MCM subunit, MCM2, and H3K9me3 as a heterochromatin marker (**Supplementary Figure S3A-F**). Finally, we calculated the ratio of the loaded MCM concentrations in heterochromatin vs euchromatin, and noted that this ratio increases from a low of ~0.7 in early G1 to ~0.9 in late G1, indicating near-equivalent origin licensing for both chromatin types by the end of G1 (**Figure 2G**).

As a control for our heterochromatin definition threshold, we randomly selected a set of 20% pixels for similar analysis. Although total MCM signal increased at all locations during G1, neither the proportion of total MCM nor the relative concentration of loaded MCM in these randomly-selected pixels changed over the course of G1 progression (**Supplementary Figure S4A, B**). Moreover, when we analyzed MCM loading with a threshold for defining heterochromatin as the 10% brightest or 50% brightest HP1 signals instead of 20%, we found the same relative dynamics, but at different absolute values (**Supplementary Figure S4C-F**). Defining heterochromatin as the 50% brightest HP1 signals without normalizing to DNA reversed the position of heterochromatin and euchromatin on the y-axis, but their increase/decrease over G1 progression remained the same, and they converged when the signals were normalized to DNA concentrations (**Supplementary Figure S4E and F**).

### The rate of MCM loading in early G1 is faster in euchromatin than in heterochromatin

Loaded MCM complexes are very stably-associated with DNA and are essentially only unloaded in S phase ^66^ or very locally displaced by active transcription ^29, 78, 79^. The remarkable stability of loaded MCM means that licensing in G1 is largely unidirectional, and the loaded MCM we detect in late G1 is the sum of all the loading that has occurred since early G1. We were therefore limited to just inferring endogenous MCM loading rates in euchromatin vs. heterochromatin rather than directly measuring the rates. To compare actual MCM loading rates in heterochromatin and euchromatin, we integrated a doxycycline-inducible mVenus-MCM3 construct into the RPE1-hTert cells with the cell cycle phase and CDK activity reporters. We induced expression for brief periods and detected a strong signal increase by both fluorescence and immunoblotting after 2 hours of induction (**Figure 3A** and **Supplementary Figure S5A**). In the absence of doxycycline, Venus-MCM3 was produced at only ~10% of the amount produced after 2 hour of induction by immunoblotting and was undetectable by microscopy (**Figure 3A and B** compare lanes 3 and 4, **Supplementary Figure S5A**, compare lanes 1 and 3). To test if the Venus-MCM3 fusion is loaded normally, we compared the dynamics of induced Venus-MCM3 with endogenous MCM3 during the cell cycle. We probed immunoblots of chromatin-enriched fractions and found that the ectopic MCM3 fusion has a similar G1 loading and S phase unloading pattern as endogenous MCM3 (**Supplementary Figure S5B**). We detected very little loaded Venus-MCM3 or endogenous MCM3 in quiescent (G0) cells as expected (**Figure 3B**, lanes 5 and 6). Thus Venus-MCM3 is a bona fide reporter for MCM loading.

**Figure 3.**
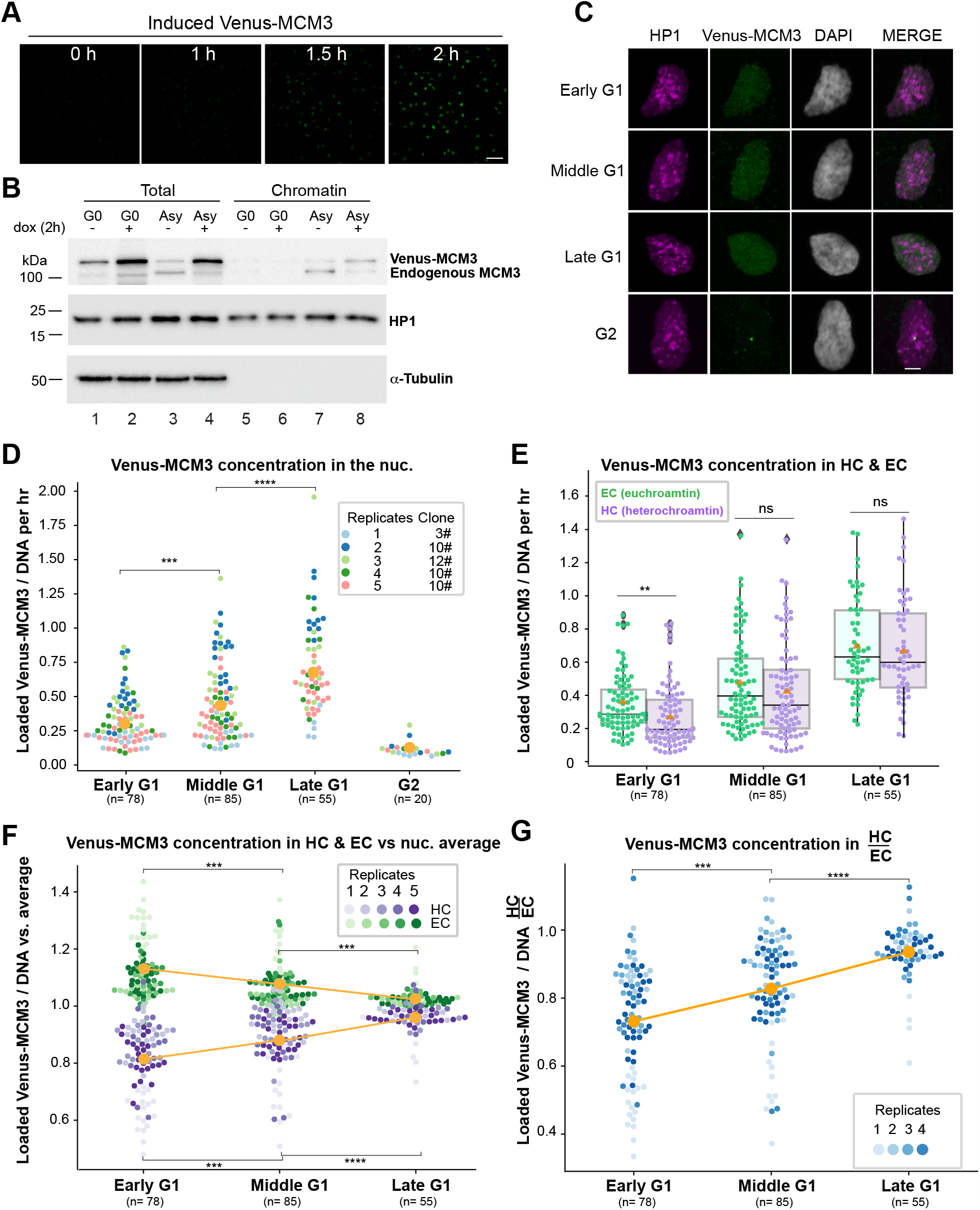
The rate of MCM loading in early G1 is faster in euchromatin than in heterochromatin. (**A**) Selected images from time-lapse imaging of RPE cells expressing doxycycline (dox)-inducible mVenus-MCM3. mVenus-MCM3 expression was recorded every 10 min after the addition 500 ng/ml dox; hours since induction is indicated, scale bar 50 μm. (**B**) Cells were made quiescent (G0) by contact inhibition or left to proliferate asynchronously (“Asy”) then treated with 500 ng/ml dox for 2 hours before harvesting. Whole cell lysates (“total”) or chromatin fractions were analyzed by immunoblotting. (**C**) Projections of 3D immunofluorescence images of representative cells after live cell imaging as in Figure 1; HP1 (magenta), Venus-MCM3 detected with anti-GFP antibody (green), DNA stained with DAPI (grey), scale bar 5 μm. Cells were treated with 500 ng/ml dox 2 hours before the end of live cell imaging. (**D, E**) Quantification of loaded mVenus-MCM3 concentration in whole nuclei (**D**) and in heterochromatin and euchromatin (**E**) after induction for 2 hours. Boxplots in **E** show median and interquartile ranges, n (number of cells) is indicated in the figure, five biological replicates are shown. P values 0.008 (early G1), 0.219 (middle G1), 0.595 (late G1). (**F**) Loaded Venus-MCM3 concentration in heterochromatin or euchromatin relative to the average loaded Venus-MCM3 concentration in whole nuclei; mean is plotted in orange. One-way ANOVA, Tukey post-hoc test, n (number of cells) is indicated in the figure. (**G**) Ratio of Venus-MCM3 concentration in heterochromatin to euchromatin; mean is plotted in orange. One-way ANOVA, Tukey post-hoc test, n (number of cells) is indicated in the figure. p value ranges are indicated as * p ≤ 0.05, ** p ≤ 0.01, *** p ≤ 0.001 and **** p ≤ 0.0001.

We then imaged asynchronously-proliferating live cells and induced Venus-MCM3 expression for the final 2 hours of imaging prior to extraction and immunostaining. In this way we measure MCM loading over a defined period of time because we restrict our analysis to the newly-synthesized Venus MCM. We detected only faint Venus fluorescence after 1 hour of induction, and almost all of the detectable Venus-MCM3 accumulated during the second hour (**Figure 3A**). We stained for loaded Venus-MCM3 using an anti-GFP antibody and HP1 as in Figure 2 and derived relative rates of Venus-MCM3 loading per hour (**Figure 3C**,**3D**). By this analysis, the overall MCM loading rate increases during G1 progression and is highest in late G1 (**Figure 3D**). We then compared relative rates in heterochromatin and euchromatin. In early G1, the MCM loading rate was significantly higher in euchromatin than in heterochromatin, and this difference disappeared gradually in middle G1 and late G1 phases (**Figure 3E**). Similar to endogenous MCM (**Figure 2F**), in early G1 Venus-MCM3 euchromatin loading was more concentrated and heterochromatin loading less concentrated than the nuclear average, but the concentrations of loaded MCM in both chromatin types were similar to each other by late G1 (**Figure 3F**). The ratio of Venus-MCM3 loading in heterochromatin to euchromatin also increased as G1 progressed **(Figure 3G**). Taken together, we conclude that the rate of MCM loading per hour increases during G1 and early G1 favors fast euchromatin loading. Strikingly, although the loading rate increases for both heterochromatin and euchromatin in middle and late G1, heterochromatin loading accelerates *more* than euchromatin does to achieve similar rates and concentrations by the end of G1.

### Global histone hyperacetylation normalizes early G1 heterochromatin and euchromatin MCM loading

We then sought to determine if manipulating chromatin in cells would cause a corresponding change in MCM loading dynamics. Since many of the largest differences between heterochromatin and euchromatin were apparent in early G1, we hypothesized that narrowing the difference between euchromatin and heterochromatin properties would promote more equal loading in early G1. To test this idea, we treated asynchronously proliferating cells for just 3 hours with increasing concentrations of the histone deacetylase inhibitor, trichostatin A (TSA), to induce histone hyperacetylation and presumably a higher proportion of open chromatin ^80^. We prioritized short treatment to reduce the confounding effects of perturbed gene expression. We then analyzed only early G1 cells as defined in **Figure 1D** and **1E**. We found that 300 nM TSA induced a strong increase in global histone H3 acetylation after 3 hours (**Figure 4A** compare lanes 1 and 4). When we analyzed overall MCM loading (all types of chromatin, throughout G1) after TSA treatment, we detected no difference between control and treated cells (**Figure 4B**). On the other hand, when we analyzed the distribution of MCM loading in early G1, we found that, as before, MCM loading favored euchromatin in control cells (ratio of HC:EC well below 1), but brief TSA treatment significantly reduced the disadvantage for heterochromatin (**Figure 4C**). These results indicate that the differences in MCM loading dynamics rely on global chromatin acetylation status.

**Figure 4.**
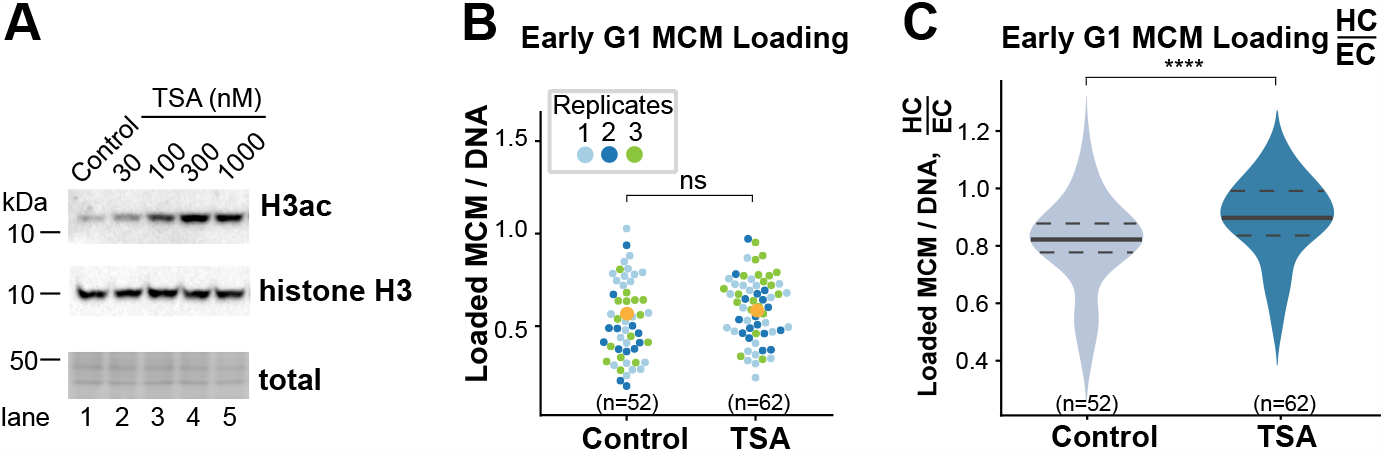
Global histone hyperacetylation normalizes early G1 heterochromatin and euchromatin MCM loading. (**A**) Representative immunoblot of histone H3 acetylation in untreated and TSA-treated asynchronously proliferating RPE1-hTert cells. Cells were treated with the indicated concentrations of TSA for 3 hours. (**B, C**) Quantification of total loaded MCM3 concentration (**B**), and the ratio of loaded MCM3 concentration in heterochromatin to euchromatin (**C**) in early G1 cells treated with 300 nM TSA for 3 hours. Three replicates are shown. One-way ANOVA test, n (number of cells) is indicated in the figure. **** p ≤ 0.0001.

### ORCA-dependent ORC loading dynamics support faster MCM heterochromatin loading

In eukaryotic cells, MCM loading to license origins starts with the DNA-binding complex, ORC, which consists of six subunits, ORC1-ORC6. Mammalian ORC selects the sites for MCM loading by mechanisms that are still incompletely understood but include interactions with chromatin features and chromatin binding proteins ^81^. ORC then cooperates with the CDC6 and CDT1 proteins to directly load MCM complexes onto DNA ^1, 2, 82^. We hypothesized that the specific MCM loading dynamics in different G1 subphases could be dictated by the distribution of loaded ORC. To test that idea, we analyzed the distribution of endogenous loaded ORC4 as a marker of the complex. The overall ORC4 concentration per unit DNA increased from early G1 to late G1 phase indicating progressive accumulation of ORC on chromatin during G1 **(Supplementary Figure S6A, B**). Interestingly and like MCM, the concentration of ORC4 per unit DNA in heterochromatin and euchromatin relative to the nuclear average was widely different in early G1 again strongly favoring euchromatin over heterochromatin, but these values converged in late G1 (**Figure 5A)**. Like endogenous MCM loading, the ratio of loaded ORC4 in heterochromatin to euchromatin increased from early to late G1 (**Figure 5B**). We observed similar trends in a stably expressed YFP-ORC1 fusion, although the effects were more modest than for endogenous ORC4 (**Supplementary Figure S6 C-F**). The ORC1 subunit can independently bind chromatin through its BAH domain ^81^, raising the possibility that the ORC1 we detect may localize to some chromatin sites independently of the full ORC. These results are consistent with a relative increase in loaded ORC concentration on heterochromatin in late G1 that could support accelerated MCM loading on heterochromatin in late G1.

**Figure 5.**
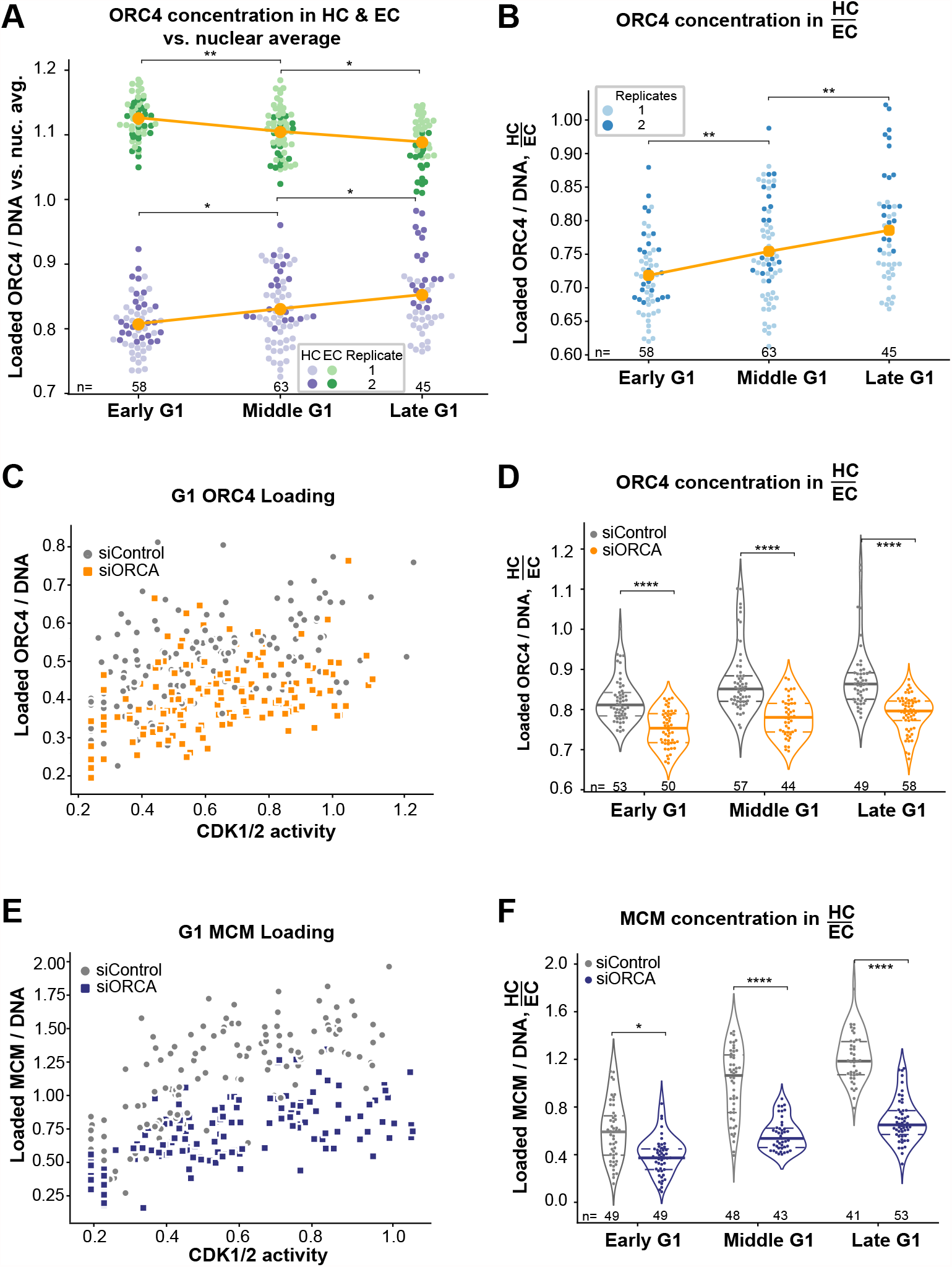
ORCA-dependent ORC loading dynamics support faster MCM heterochromatin loading. (**A**) Loaded ORC4 concentration (ORC4 signal per unit DAPI) in heterochromatin (purple) and euchromatin (green) relative to the overall loaded ORC4 concentration in each nucleus; mean is plotted in orange. Two replicates are shown. (**B**) Ratio of loaded ORC4 concentration in heterochromatin to euchromatin; mean is plotted in orange. Two replicates are shown. One-way ANOVA, Tukey post-hoc test, n (number of cells) is indicated in the figure. (**C, D**) ORC4 concentration (**C**) and the ratio of ORC4 concentration in heterochromatin to euchromatin pixels (**D**) in G1 cells treated with 100 nM control siRNA or ORCA siRNA for 48 hours. Two biological replicates are shown. One-way ANOVA, Tukey post-hoc test, n (number of cells) is indicated in the figure. (**E, F**) MCM3 concentration (**E**) and the ratio of endogenous MCM3 concentration in heterochromatin to euchromatin (**F**) in G1 cells treated with 100 nM control siRNA or ORCA siRNA for 48 hours. Two biological replicates were shown. One-way ANOVA, Tukey post-hoc test, n (number of cells) is indicated in the figure. In all panels p value ranges are indicated as * p ≤ 0.05, ** p ≤ 0.01, *** p ≤ 0.001 and **** p ≤ 0.0001.

To understand how ORC loading dynamics change during G1, we turned our attention to the role of the ORCA/LRWD1 protein. ORCA interacts with both ORC and heterochromatin marks, and ORCA plays a role in both ORC recruitment to heterochromatin loci and also heterochromatin organization ^40, 83^. We considered that ORCA may be required for the dynamics of both ORC and MCM loading as G1 progresses. We used RNAi to reduce ORCA and observed the previously reported reduction in ORC chromatin association and the heterochromatin mark histone H3 lysine nine trimethylation (H3K9me3) (**Supplementary Figure 7A**, compare lanes 3 and 4 ^41, 84^). ORCA depletion for 72 hours caused a moderate increase in S phase cells **(Supplementary Figure 7B)**. As measured by analytical flow cytometry of cells extracted to remove soluble proteins, ORCA depletion also specifically decreased the level of chromatin-loaded ORC4 in G1 phase cells (**Supplementary Figure 7C-D**). By quantitative immunofluorescence, ORCA depletion reduced the concentrations of both ORC4 (**Figure 5C**) and MCM (**Figure 5E**) on DNA in all G1 subphases (**Supplementary Figure 7E and F**). Importantly, ORCA depletion disrupted not only the total increase of ORC4 loading, but also the normal relative increase in the ratio of heterochromatin to euchromatin ORC4 loading (**Figure 5D**). Correspondingly, ORCA was also required for the relative increase in MCM loading on heterochromatin vs. euchromatin during G1 progression (**Figure 5F**).

### Preferential heterochromatin under-licensing in shortened G1 cells

Because we established that MCM loading in euchromatin is advanced relative to heterochromatin in early G1 (**Figure 2**), we hypothesized that a premature G1/S transition will have a preferential negative impact on heterochromatin replication. Our reasoning is that an early G1/S transition could occur when euchromatin is more fully licensed whereas heterochromatin licensing is less complete. To test this hypothesis, we expressed human cyclin E from a doxycycline-inducible promoter. Cyclin E normally accumulates in late G1, where it promotes S phase entry and progression by activating CDK2 ^85, 86^. We and others had previously shown that cyclin E overproduction shortens G1 and can cause cells to enter S phase with less total loaded MCM (**Supplementary Figure S8A, B** ^54, 87, 88^). Treating cells with doxycycline for 7 hours strongly induced cyclin E and significantly shortened G1 length from a mean of 7 hours to 3 hours (**Figure 6 A, B**). Under these conditions, cyclin E overexpression also induced high CDK1/2 activity at much younger physical ages (**Figure 6C**). In these cells the overall concentration of loaded MCM achieved within these shorter G1 phases was also clearly less than in control cells (**Figure 6D, E**). This under-licensing was more profound in heterochromatin than euchromatin especially in middle and late G1 cells (**Figure 6E**). Interestingly, when we compared euchromatin and heterochromatin, we found that heterochromatin MCM loading didn’t reach the same concentration as euchromatin loading even in late G1 cells, whereas in the control cells both types of chromatin had similar MCM loading in late G1 as before (**Figure 6F**). Since we had manipulated CDK1/2 activity by cyclin E overexpression, we categorized G1 subphases by their physical age rather than molecular age for cyclin E-overproducing cells as indicated on the x-axis of **Figure 6C**. Compared to control G1 cells, loaded MCM distribution in cyclin E-overproducing cells favored euchromatin throughout all G1 subphases, and the two chromatin types never reached parity. These results support the conclusion that a premature G1/S transition preferentially causes underlicensed heterochromatin.

**Figure 6.**
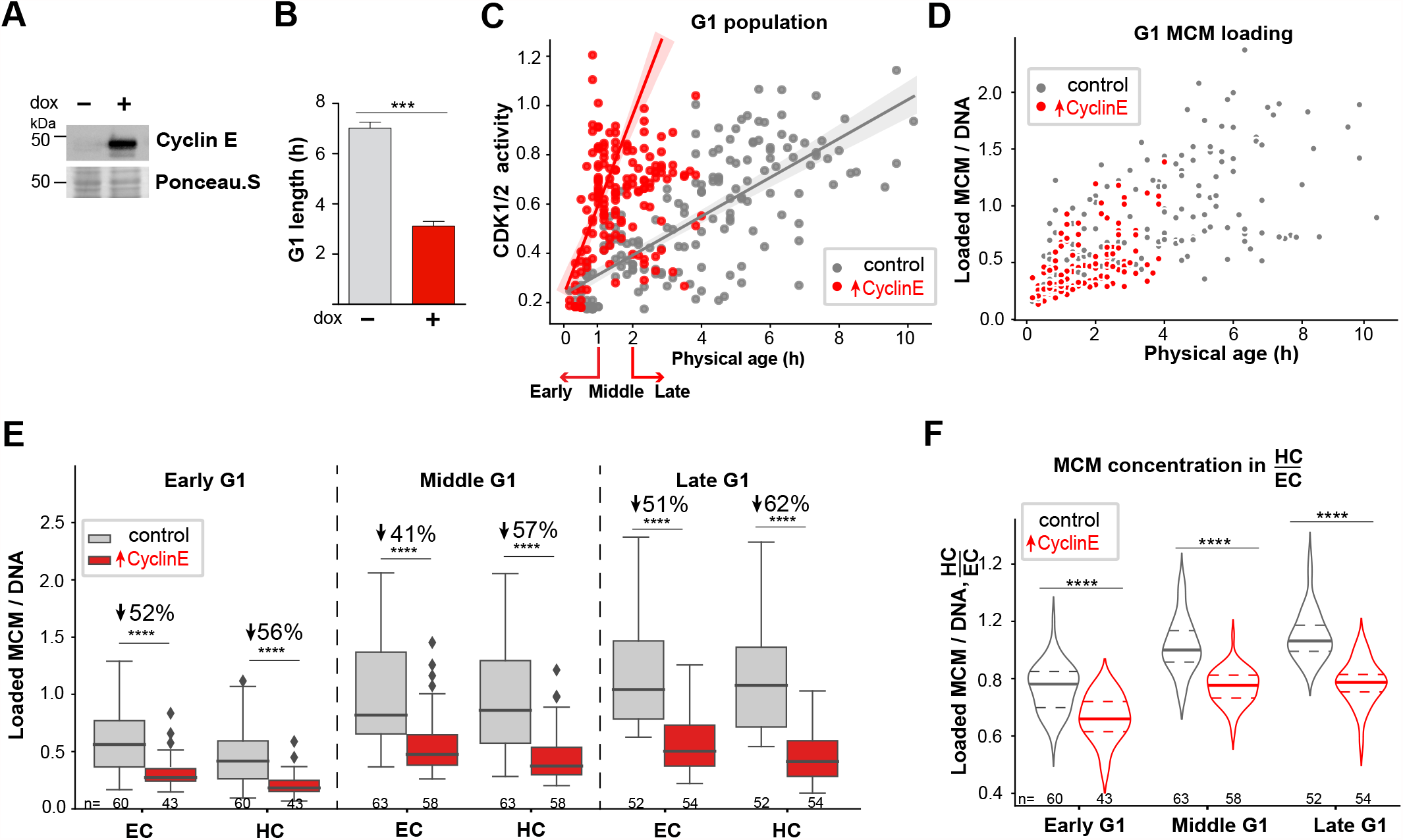
Preferential heterochromatin under-licensing in shortened G1 cells. (**A**) Ectopic cyclin E expression was induced by treatment with 15 ng/ml doxycycline (dox) for 7 hours prior to analysis by immunoblotting. (**B**) G1 length of cells in **A** measured by live cell imaging of the PCNA S phase reporter. T-test, n (number of cell) =40 for each group. (**C**) CDK1/2 activity relative to physical age in control (grey) or cyclin E-expressing (red) G1 phase cells treated as in **B**. (**D**) Loaded MCM3 concentration relative to physical age in control (grey) or cyclin E-expressing (red) G1 cells as in **B**. (**E**) Loaded MCM concentration in heterochromatin and euchromatin in G1 subphases in control or cyclin E-expressing cells. G1 subphases for E and F were defined for control cells as in Figure 1 and for cyclin E-overproducing cells by physical age as indicated in **B**. Boxplots show median and interquartile ranges, one-way ANOVA, Tukey post-hoc test, n (number of cells) is indicated in the figure. (**F**) Quantification of the ratio of loaded MCM concentration in heterochromatin to euchromatin in control or cyclin E-expressing G1 cells. Violin plots indicate the median and interquartile ranges, one-way ANOVA, Tukey post-hoc test, n (number of cells) is indicated in the figure. In all panels p value ranges are indicated as * p ≤ 0.05, ** p ≤ 0.01, *** p ≤ 0.001 and **** p ≤ 0.0001.

### Heterochromatin is more vulnerable than euchromatin to under-replication and DNA damage

Reduced origin licensing in G1 phase could leave segments of DNA under-replicated in S phase, thus threatening genome stability. Based on our findings that heterochromatin MCM loading is relatively slower in G1 phase, we postulated that the consequences of a premature G1/S transition include preferential heterochromatin under-replication during the following S phase resulting in more DNA damage in heterochromatin. We therefore examined the recruitment of p53 binding-protein 1 (53BP1), a biomarker for local replication stress and DNA double-strand breaks ^89, 90^ to different chromatin types in very late S phase immediately after cyclin E induction. We first analyzed overall 53BP1 loading per nucleus in control and cyclin E-overproducing cells (similar to our analysis of MCM and ORC loading) and detected no differences between the two groups (**Figure 7A, B**, untreated and “↑Cyclin E”). We also treated control and cyclin E-overproducing cells briefly (4 hours) with a low dose (25 ng/ml) of aphidicolin, a DNA polymerase inhibitor that induces replication stress and DNA damage. Under these conditions, aphidicolin treatment did not induce significant global DNA damage in control cells, but it did modestly increase overall DNA damage in cyclin E-overproducing cells as measured by 53BP1 recruitment (**Figure 7A, B**, *aphidicolin* compared to *↑Cyclin E + aphidicolin*).

**Figure 7.**
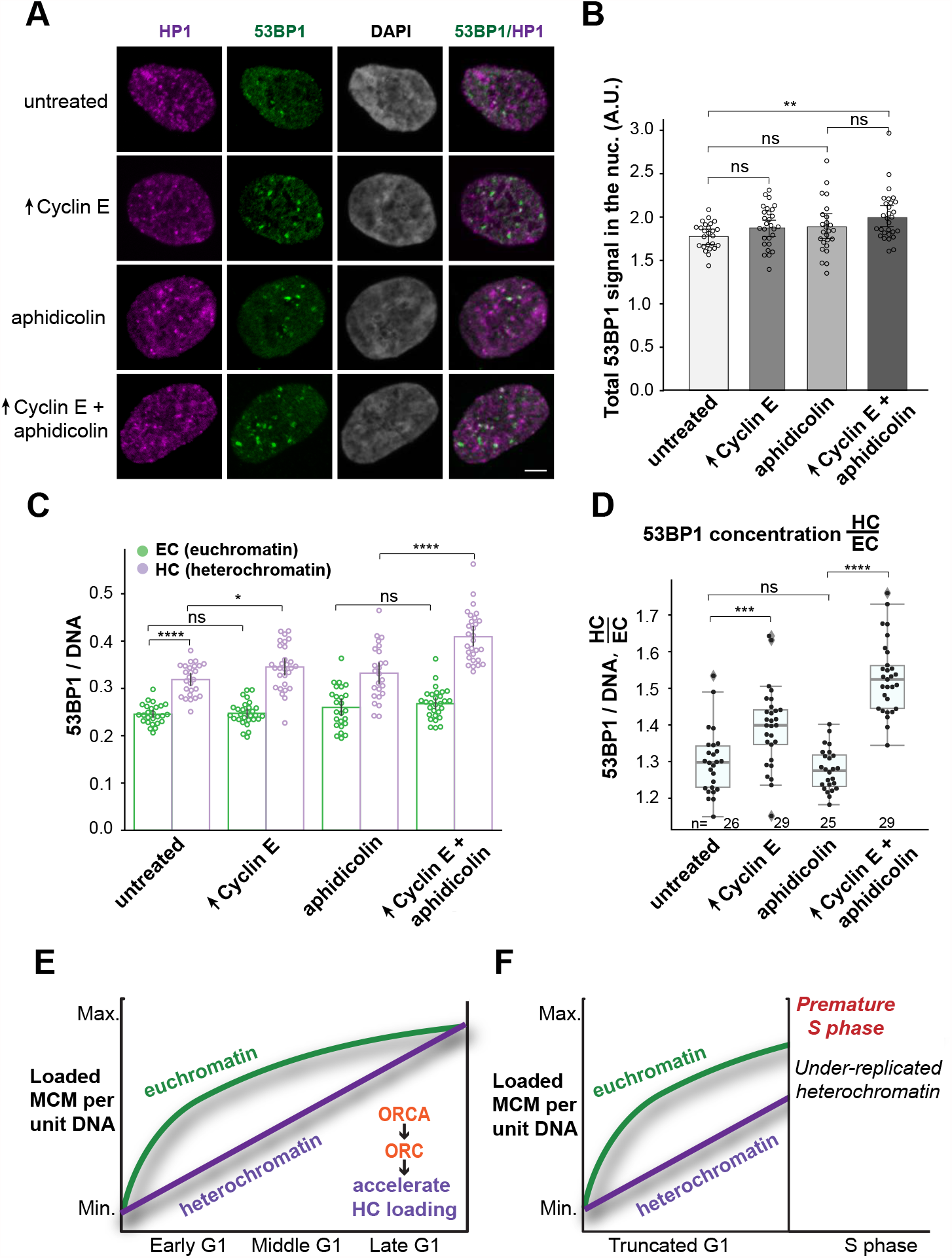
Heterochromatin is more vulnerable than euchromatin to under-replication and DNA damage. (**A**) Projections of 3D immunofluorescence images of representative late S phase cells after live cell imaging as in Figure 1. Cells were treated with 500 ng/ml dox for 18 hours before imaging and with vehicle or 25 ng/ml aphidicolin for the final 4 hours of imaging as indicated. Soluble proteins were extracted prior to fixation and immunostaining for endogenous 53BP1 (green), HP1 (magenta), and DNA (DAPI, grey) scale bar 5 μm. Cells with G1 phase shorter than 4 hours were selected for analysis in the next late S phase using PCNA variance as an indicator of S phase progression. (**B, C**) Quantification of total bound 53BP1 signal in late S phase nuclei (**B**) and (**C**) the concentration of 53BP1 signal in heterochromatin (purple) or euchromatin (green). (**D**) Ratio of 53BP1 concentration in heterochromatin to euchromatin; n (number of cells) is indicated in the figure. Two replicates are shown. In all panels p value ranges are indicated as * p ≤ 0.05, ** p ≤ 0.01, *** p ≤ 0.001 and **** p ≤ 0.0001. (**E**) Illustration of normal MCM loading dynamics in euchromatin vs heterochromatin. (**F**) Illustration of heterochromatin under-licensing from premature G1/S transition. See Discussion for details.

To explore the intranuclear DNA damage distribution between the different chromatin types, we analyzed the location of 53BP1 signals in heterochromatin and euchromatin. We noted that even in untreated cells, the concentration of 53BP1 in heterochromatin was higher than in euchromatin (**Figure 7C**). Cyclin E overproduction for 18 hours (less than one full RPE1-hTert cell cycle) caused even more 53BP1 recruitment to heterochromatin, but strikingly, had no effect on euchromatin (**Figure 7C**). This effect was exacerbated by low-dose aphidicolin treatment for 4 hours (**Figure 7C)**. As we did for MCM distribution, we calculated the distribution of 53BP1 concentrations. 53BP1 recruitment induced after cyclin E overproduction occurred preferentially in heterochromatin, and this difference was exacerbated by aphidicolin treatment (**Figure 7D**). Furthermore, we analyzed the chromatin association and focal localization of the single-stranded DNA binding protein, replication protein A (RPA), specifically in the first G2 phase following cyclin E induction and/or aphidicolin treatment. Notably, cyclin E-overproducing cells accumulated many RPA foci in G2 phase, suggesting the persistence or exposure of ssDNA from incomplete replication (**Supplementary Figure S9A-C**). Aphidicolin alone did not induce G2 phase RPA foci but did enhance the effects of cyclin E overproduction. Overall RPA signal was also preferentially distributed to heterochromatin over euchromatin (**Supplementary Figure S9D and E**). Altogether, these data strongly suggest that precocious S phase entry is particularly detrimental for heterochromatin replication, and that this disadvantage for heterochromatin is at least partly because these regions are more under-licensed than euchromatic regions (**Figure 7E and F**). Preferential under-licensing in heterochromatin is itself attributable to the unique dynamics of MCM loading in heterochromatin compared to euchromatin within G1 phase.

## DISCUSSION

In this study, we combined time-lapse imaging with fixed cell immunostaining to define the dynamics of human MCM loading *within* G1 phase. To our knowledge this is the first quantitative analysis of MCM loading in single cells at such high temporal resolution with respect to both physical age (in hours since cell division) and also “molecular age” defined by CDK2 activity. We note that the dynamics of MCM loading can be analyzed relative to any marker for which there are suitable detection reagents. Identifying where and how MCM is loaded within distinct genomic regions in G1 typically poses technical challenges because MCM is at all replication forks during S phase. Moreover, as in most eukaryotes, human origins are highly flexible and variable between individual cells which reduces the information obtained from analyzing cell populations. We instead developed methods to quantify relative rates of MCM loading in single G1 cells by comparing two primary genomic environments, heterochromatin and euchromatin. This strategy also avoids the potential pitfalls of artificial cell cycle synchronization or the averaging effect of analyzing bulk cell populations. Here we report the first comprehensive analysis of G1 MCM loading dynamics in different intranuclear environments, the mechanism driving those dynamics, and the downstream consequences those dynamics can have.

We uncovered a preference for MCM loading in euchromatin, but surprisingly, this preference is largely confined to cells in early G1 phase. Based on the long history of the transcriptional regulation field, it is intuitive that the more accessible euchromatin regions have an advantage over less accessible heterochromatin. MCM loading may be easier or simpler in a more accessible chromatin environment, and thus faster. What was *not* anticipated at the outset of this study is that the rate of heterochromatin loading increases relative to euchromatin loading during middle and late G1 subphases. As a result, heterochromatin loading lags behind euchromatin in early G1, but differences in acceleration allows heterochromatin to “catch up” with euchromatin by the end of G1. Other studies of MCM binding sites in mammalian cells also report similar overall loading in both transcriptionally active and repressed genomic loci which generally correlate with euchromatin and heterochromatin respectively ^16, 29^.

Thus far, attention to the differences between the two chromatin types in DNA replication has been almost exclusively focused on the timing of origin firing within S phase itself. Origin firing time is a combined effect of origin licensing in G1 to define all potential origins and the recruitment of origin initiation factors in a process that is also influenced by chromatin ^40, 42, 91^. In many species, replication timing in S phase shows a general (though not exclusive) pattern of early euchromatin replication and late heterochromatin replication ^92, 93^. Differences in the density of G1 MCM loading at different individual loci have been implicated in these S phase replication timing differences; more ORC or MCM loading in G1 correlates with early firing origins ^29, 31, 94^. Early firing in S and early G1 origin licensing may both be consequences of general DNA accessibility in euchromatin. We also speculate that the higher density of MCM loading in early-replicating regions is partly from loading very quickly early in G1 and then having the entire rest of G1 phase to add additional MCM complexes at the same sites.

The molecular mechanism driving differential loading rates correlates with differences in ORC chromatin binding, and the ORC and MCM chromatin binding differences themselves require the ORCA/LRWD1 protein. ORCA plays multiple roles in chromosome biology, including ORC recruitment to heterochromatin through a direct interaction with H3K9me3 ^84^. Thus only some genomic regions (heterochromatin) rely on help from ORCA-mediated ORC recruitment. We envision that accessible euchromatic regions recruit ORC for MCM loading without much assistance, but less accessible regions require helper factors. These additional factors may be specialized for different types of chromatin or may cooperate with one another to ensure MCM loading in even the most inaccessible regions. For example, trimethylation of histone H4 lysine 20 has also been implicated in heterochromatin licensing at a subset of ORCA-bound sites ^95^. These mechanisms enhance MCM loading in heterochromatin regions, but equitable licensing distribution can also be promoted by factors that suppress MCM loading in euchromatin. Two recent studies of yeast origins described mechanisms to reduce disparities in MCM loading levels among different origins ^42, 96^; analogous mechanisms may also operate in mammalian cells. We are also intrigued by the apparent slowing of euchromatin loading acceleration between early G1 and later G1 times. It is possible that a passive mechanism improves MCM loading in heterochromatin later in G1 phase because the more accessible euchromatin sites are already occupied ^97–99^.

Because heterochromatin reaches full licensing closer to the end of G1 phase than euchromatin does, heterochromatin is more vulnerable to any change that causes premature S phase entry (**Figure 7E**). Early S phase entry may happen stochastically from random fluctuations in gene expression leading to early cyclin E/ CDK2 activation, or it may happen chronically if cells acquire genetic or epigenetic alterations that shorten G1 phase, such as oncogene activation or tumor suppressor loss. Importantly, it is the rate of licensing combined with the length of G1 phase that determines how much overall under-licensing cells experience, and the relative rates in different regions determine where under-licensing will be most severe. We note that cyclin E overproduction truncated G1 without affecting the apparent MCM loading rate (**Figure 6D**). We demonstrate that euchromatin is also somewhat under-licensed in cells with artificially short G1 phases, but not to the extent that it increases genome damage in euchromatin. On the other hand, heterochromatin is much more underlicensed and the increase in under-replication and DNA damage was largely confined to heterochromatin (**Figure 7F**). This concentration of under-replication in heterochromatin is presumably from both under-licensing in G1 and late origin firing in S phase which leaves even less time for replication to finish ^92, 93^. Highly compacted heterochromatin could also be a barrier for timely DNA repair factor recruitment ^100–102^. Interestingly, a period of cyclin E overproduction in a prior report caused some large genomic deletions ^103^. In that study, those deleted sequences were primarily late-replicating (13 of 16 deletions analyzed) and, based on replication timing, we presume are associated with heterochromatin and delayed origin licensing. Our results suggest that the final stage of G1 is crucial for heterochromatin to become fully licensed and therefore fully replicated.

Finally, we note that the distribution of euchromatin and heterochromatin varies by cell type. We predict that differences in facultative heterochromatin that distinguish one cell type from another are also among the regions most vulnerable to late origin licensing and under-replication in those cell types. We also presume that constitutive heterochromatin is hypersensitive to under-licensing in most cell types, but that notion remains to be explored. Licensing dynamics may be unique in centromeres, telomeres, or other distinct chromatin subregions, and additional investigations using other localization markers can reveal those dynamics. We also note that both chromatin structure and G1 length are altered in many cancers ^104–106^. The insights gained from this study can contribute to understanding both the source and location of genome instability in cells with such perturbations.

## Supporting information

Mei Supp video

Mei et al Supplemental Figures

## AVAILABILITY

GitHub repository (https://github.com/purvislab/MCM_project)

## ACKNOWLEDGEMENTS

We thank Supriya Prasanth and Sabrina Spencer for the generous gifts of antibodies and reagents, and we thank Abid Khan, Robert Duronio, Cook lab members, and colleagues in the field for discussion and comments on the manuscript. We thank Jeffrey Jones for research support assistance.

## Supplementary video

Representative 3D immunofluorescence image of a single asynchronously proliferating RPE1-hTert cell in G1 phase. Detergent extraction was performed before fixation as in Figure 1. Cells were stained with MCM3 (green) antibody, HP1 (magenta) antibody, and with DAPI for DNA (blue).

**Supplementary Figure 1. Specificity of MCM3 and heterochromatin detection**. (A) Immunofluorescence of asynchronously proliferating RPE1-hTert cells treated with control or MCM3 siRNAs at 100 nM for 48 hours. Detergent extraction was performed before fixation as in Figure 1. (B) Immunoblot of whole protein lysates of cells treated as in A. (C) Immunofluorescence of RPE1-hTert cells extracted and stained with MCM3 and MCM2 antibodies. (D) Immunofluorescence of RPE1-hTert cells transfected with YFP tagged ORC1 or YFP-empty vector (EV). Cells were processed as in C staining for MCM3 and GFP. (E, F) Immunofluorescence of RPE1-hTert cells stained with acetylated histone H4 (H4ac) and HP1 antibodies; plot profile of the immunofluorescence intensity signals along the yellow line through the nucleus in the boxed region of interest. (G, H) Immunofluorescence of RPE1-hTert cells with antibodies to histone H3 lysine nine trimethylation (H3K9Me3) and HP1; plot profile of the immunofluorescence intensity signals along the yellow line through the nucleus in the boxed region of interest. Scale bars are indicated in the figure panels.

**Supplementary Figure 2. MCM3 and HP1 levels on chromatin during G1 phase**. (A) Last frame of live cell imaging showing the CDK1/2 reporter localization (a), followed by immunofluorescence staining of loaded MCM using the MCM3 antibody and DAPI (b). Cells marked by arrows are sister cells of the same physical age but different “molecular ages” as defined by CDK activity; scale bar 15 μm. (B) Total Loaded MCM3 signal in the nucleus grouped by G1 subphase defined in Figure 1. (C, D) Endogenous total loaded HP1 and the ratio of HP1/DAPI in heterochromatin to the average in the nucleus. (E) DAPI intensity per unit volume in heterochromatin and euchromatin defined by HP1 immunostaining. (F) Loaded MCM3 signal from C normalized to DAPI signal and grouped by G1 subphase. One-way ANOVA, Tukey post-hoc test, n (number of cells) is indicated in the figure. In all panels p value ranges are indicated as * p ≤ 0.05, ** p ≤ 0.01, *** p ≤ 0.001 and **** p ≤ 0.0001.

**Supplementary Figure 3. Differential dynamics of MCM2 loading in euchromatin and heterochromatin (similar to MCM3 analysis in Figure 2)**. (A) Projections of 3D immunofluorescence images of representative cells after live cell imaging as in Figure 1; H3K9me3 (magenta), endogenous MCM2 (green), scale bar represents 5 µm. (B, C) Total loaded MCM2 signal relative to CDK1/2 activity in G1 or G2 cells; cells in C are grouped by G1 subphase. (D) Loaded MCM2 normalized to DAPI in G1 subphases. (E) Proportion of total loaded MCM2 in heterochromatin (purple) and euchromatin (green) in early, middle, and late G1 cells. (F) Loaded MCM2 concentration in heterochromatin (purple) or in euchromatin (green) relative to average loaded MCM2 in nuclei; means are plotted in orange. One-way ANOVA, Tukey post-hoc test, n (number of cells) is indicated in the figure. In all panels p value ranges are indicated as * p ≤ 0.05, ** p ≤ 0.01, *** p ≤ 0.001 and **** p ≤ 0.0001.

**Supplementary Figure 4. Controls for defining heterochromatin in immunofluorescence images**. (A) Loaded MCM3 in a random 20% selection of pixels relative to CDK1/2 activity in G1 cells plotted as the proportion of total MCM signal in nuclei. (B) Loaded MCM3 concentration in the random 20% selection of pixels in A relative to average nuclear loaded MCM3 concentration. (C-F) Quantification of the proportion of loaded MCM3 (C and E) or the concentration relative to the nuclear average (D and F) for heterochromatin defined as the 10% or 50% brightest HP1 signals in the indicated G1 subphases. Mean is plotted in orange; one-way ANOVA, Tukey post-hoc test, n (number of cells) is indicated in the figure. In all panels p value ranges are indicated as * p ≤ 0.05, ** p ≤ 0.01, *** p ≤ 0.001 and **** p ≤ 0.0001.

**Supplementary Figure 5. Venus-MCM3 is loaded onto chromatin similarly to endogenous MCM3**. (A) Immunoblot of total protein lysates of asynchronously proliferating RPE1-hTert cells with stably-integrated inducible mVenus-MCM3 treated with 500 ng/ml doxycycline (dox) for 0, 1, 2, 3, or 6 hours. (B) Asynchronous cells or cells synchronized by contact-inhibition (G0) and release were pre-treated with 500 ng/ml dox for 24 hours. Cells were harvested after release at 16, 18, 21, 24 and 30 hours to enrich for G1, late G1/early S, mid-S phase, late S and G2 phase respectively. Whole-cell lysates and chromatin fractions were probed for the indicated proteins.

**Supplementary Figure 6. The concentration of loaded ORC increases during G1 phase**. (A) Projections of 3D immunofluorescence images of representative cells after live cell imaging as in Figure 1. Loaded endogenous ORC4 (green) and HP1 (magenta) were detected by immunostaining; scale bar represents 5 µm. (B) Loaded ORC4 concentration normalized to DNA in G1 subphases; two biological replicates are shown; Means are plotted in orange; n (number of cells) is indicated in the figure. One-way ANOVA, Tukey post-hoc test, n (number of cells) is indicated in the figure. (C) As in A except that cells were transfected with plasmid to express dox-inducible YFP-ORC1, treated with 100 ng/ml dox for 48 hours, then extracted, fixed, and stained with GFP antibody; HP1 (magenta), YFP-ORC1 (green), scale bar 5 μm. (D) Quantification of YFP-ORC1 concentration in subphases of G1; two biological replicates are shown; Means are plotted in orange. One-way ANOVA, Tukey post-hoc test, n (number of cells) is indicated in the figure. (E) Loaded YFP-ORC1 concentration (YFP signal per unit DAPI) in heterochromatin (purple) and euchromatin (green) relative to the nuclear average loaded YFP-ORC1 signal; two replicates are shown. One-way ANOVA, Tukey post-hoc test, n (number of cells) is indicated in the figure. (F) Ratio of ORC4 concentration in heterochromatin to euchromatin; two replicates are shown. One-way ANOVA, Tukey post-hoc test, n (number of cells) is indicated in the figure. In all panels p value ranges are indicated as * p ≤ 0.05, ** p ≤ 0.01, *** p ≤ 0.001 and **** p ≤ 0.0001.

**Supplementary Figure 7. ORCA depletion impairs ORC4 loading and MCM loading**. (A) Asynchronously proliferating RPE1-HTert cells were transfected with 100 nM siRNA targeting ORCA or control siRNA for 72 hours before harvesting. Samples were fractionated into DNA-loaded chromatin fractions and total lysate for immunoblotting for the indicated proteins. (B, C, D) Flow cytometry of chromatin-bound proteins in EdU-labeled cells treated as in A. The distribution of cells in cell cycle phases (B) and loaded ORC4 (C) is shown. Quantification of loaded ORC4 in the G1 gate (rectangles in C) is shown in (D). (E, F) Cells were treated with control siRNA or siRNA targeting ORCA for 48 hours. Loaded ORC4 (E) and MCM3 (F) immunostaining signals were normalized to DAPI. One-way ANOVA, Tukey post-hoc test, n (number of cells) is indicated in the figure. In all panels p value ranges are indicated as * p ≤ 0.05, ** p ≤ 0.01, *** p ≤ 0.001 and **** p ≤ 0.0001.

**Supplementary Figure 8. Cyclin E overexpression shortens G1 and induces under-licensing**. (A, B) Flow cytometry of chromatin-bound proteins in EdU-labelled cells treated with 100 ng/ml dox to induce ectopic cyclin E expression for 24 hours. The distribution of cell cycle phases (A) and loaded MCM (B) are shown. Replicate results shown in Matson et al. 2017.

**Supplementary Figure 9. Heterochromatin is more vulnerable than euchromatin to under-replication and DNA damage (similar to 53BP1 analysis in Figure 7)** (**A**) Projections of 3D immunofluorescence images of representative G2 phase cells after live cell imaging as in Figure 1. Cells were treated with 500 ng/ml dox for 18 hours before imaging and with vehicle or 25 ng/ml aphidicolin for the final 4 hours of imaging as indicated. Soluble proteins were extracted prior to fixation and immunostaining for endogenous RPA (green), HP1 (magenta), and DNA (DAPI, grey) scale bar 5 μm. Cells with G1 phases shorter than 4 hours were selected for analysis in the next G2 phase using PCNA variance as an indicator of the S/G2 transition. (**B, C**) Quantification of total loaded RPA signal in G2 phase nuclei (**B**) and the number of RPA foci per cell (**C**). (**D**) Proportion of RPA in heterochromatin. (**E**) Ratio of RPA concentrations in heterochromatin to euchromatin; n (number of cells) >50 for each group; two replicates are shown. In all panels p value ranges are indicated as * p ≤ 0.05, ** p ≤ 0.01, *** p ≤ 0.001 and **** p ≤ 0.0001.

