## Supplementary material for "The consequences of differential origin licensing dynamics in distinct chromatin environments": Mei et al Supplemental Figures

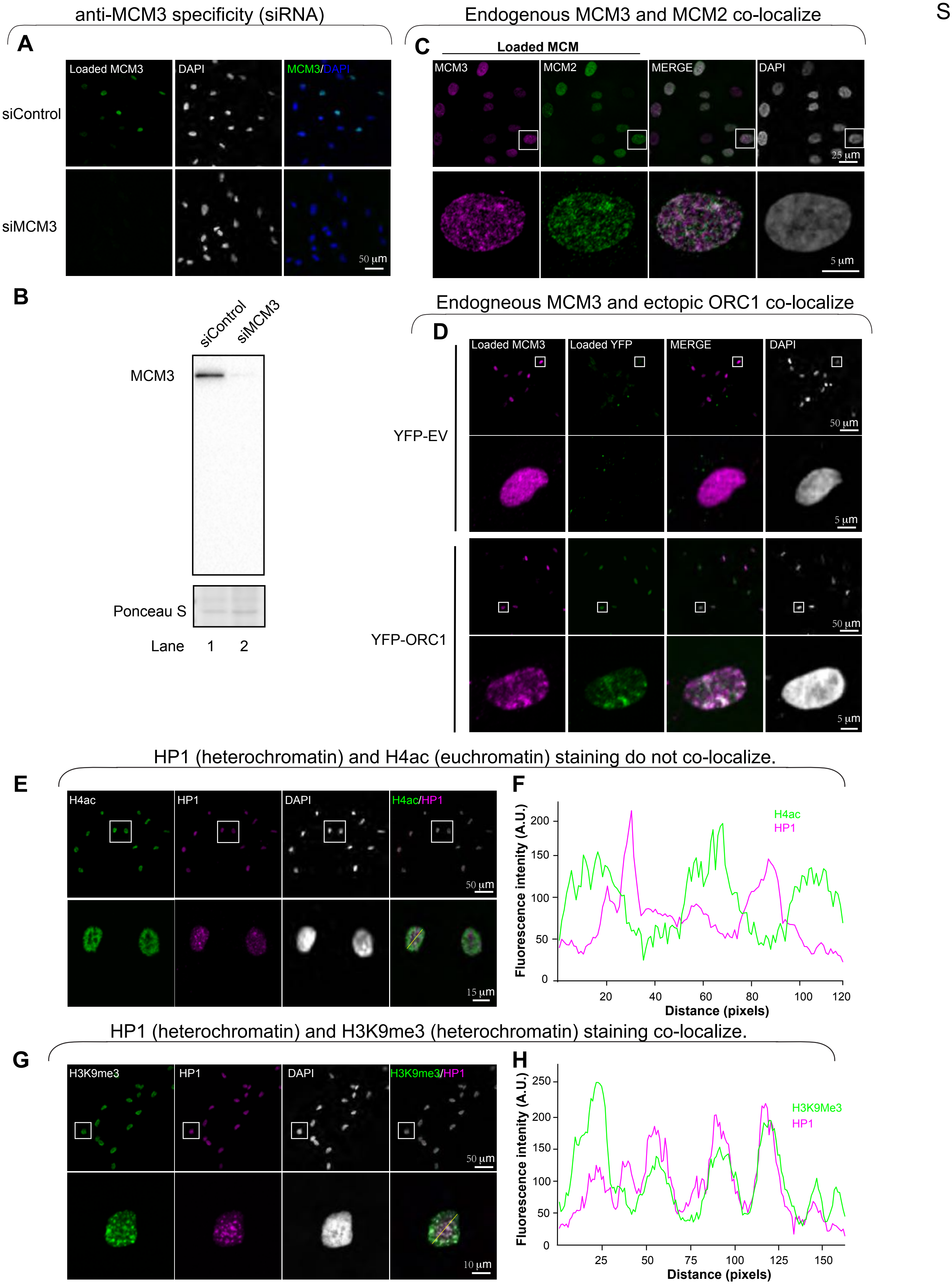


**Supplementary Figure 1. Specificity of MCM3 and heterochromatin detection.**
(**A**) Immunofluorescence of asynchronously proliferating RPE1-hTert cells treated with control or MCM3 siRNAs at 100 nM for 48 hours. Detergent and salt extraction and fixation as in Figure 1.
(**B**) Immunoblot of whole protein lysates of cells treated as in **A**.
(**C**) Immunofluorescence of RPE1-hTert cells. Cells were extracted and stained with MCM3 and MCM2 antibodies.
(**D**) Immunofluorescence of RPE1- hTert cells transfected with YFP tagged ORC1or YFP-empty vector (EV). Cells were processed as in **C** staining for MCM3 and GFP.
(**E, F**) Immunofluorescence of RPE1- hTert cells stained with acetylated histone H4 (H4ac) and HP1 antibodies; plot profile of the immunofluorescence intensity signals along the yellow line through the nucleus in the boxed region of interest.
(**G, H**) Immunofluorescence of RPE1- hTert cells with antibodies to histone H3 lysine nine trimethylation (H3K9Me3) and HP1; plot profile of the immunofluorescence intensity signals along the yellow line through the nucleus in the boxed region of interest. Scale bars are indicated in the figure panels.

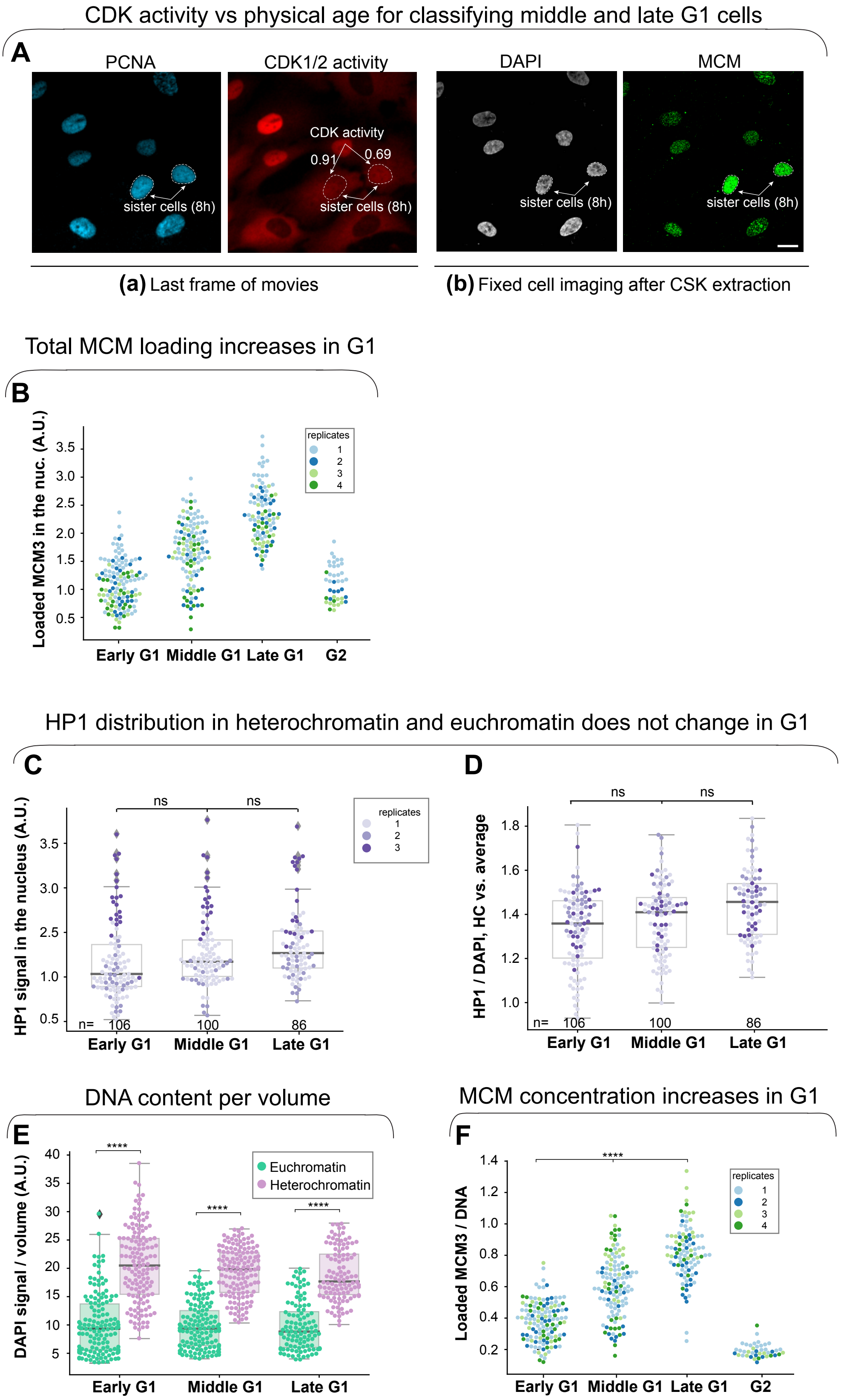


**Supplementary Figure 2. MCM3 and HP1 levels on chromatin during G1 phase.**
(**A**) Last frame of live cell imaging showing the CDK1/2 reporter localization (a), followed by immunofluorescence staining of loaded MCM using the MCM3 antibody and DAPI (b). Cells marked by arrows are sister cells of the same physical age but different "molecular ages" as defined by CDK activity; scale bar 15 μm.
(**B**) Total Loaded MCM3 signal in the nucleus relative to CDK1/2 activity; cells are color-coded by G1 subphase defined in Figure 1.
(**C, D**) Endogenous total loaded HP1 and the ratio of HP1/DNA in heterochromatin to the average in the nucleus.
(**E**) DAPI signal per unit volume in euchroamtin and in heterochromatin defined by HP1 immunostaining.
(**F**) Loaded MCM3 signal from **C** normalized to DAPI signal and grouped by G1 subphase. One-way ANOVA, Tukey post-hoc test, n (number of cells) is indicated in the figure. In all panels p value ranges are indicated as * $p \leq 0.05$, ** $p \leq 0.01$, *** $p \leq 0.001$ and **** $p \leq 0.0001$.

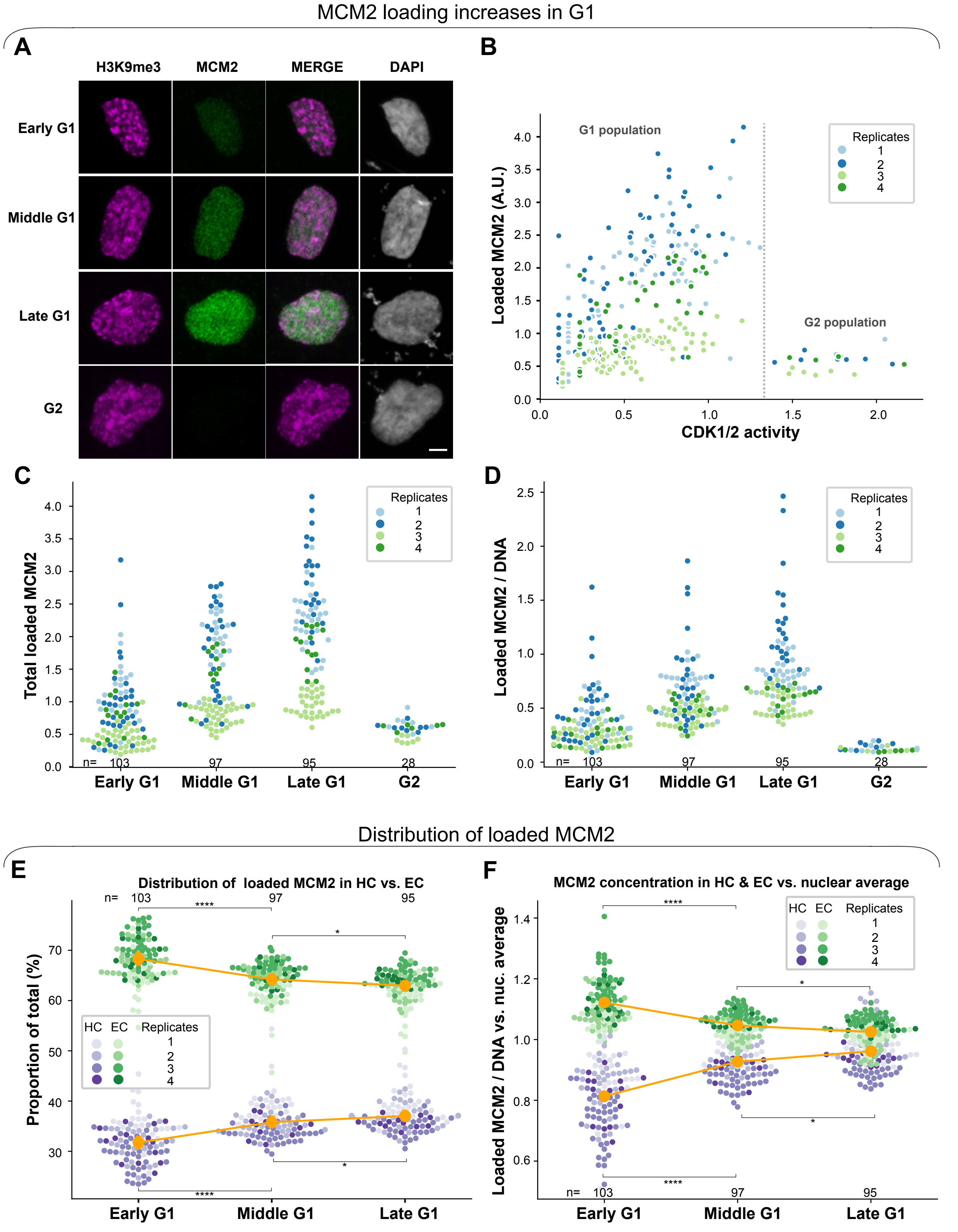


**Supplementary Figure 3. Differential dynamics of MCM2 loading in euchromatin and heterochromatin** (**similar to MCM3 analysis in Figure 2**).

(**A**) Projections of 3D immunofluorescence images of representative cells after live cell imaging as in Figure 1; H3K9me3 (magenta), endogenous MCM2 (green), scale bar represents 5 µm.

(**B, C**) Total loaded MCM2 signal relative to CDK1/2 activity in G1 or G2 cells; cells in **C** are grouped by G1 subphase.

(**D**) Loaded MCM2 normalized to DAPI in G1 subphases.

(**E**) Proportion of total loaded MCM2 in heterochromatin (purple) and euchromatin (green) in early, middle, and late G1 cells.

(**F**) Loaded MCM2 concentration in heterochromatin (purple) or in euchromatin (green) relative to average loaded MCM2 in nuclei; means are plotted in orange. One-way ANOVA, Tukey post-hoc test, n (number of cells) is indicated in the figure.

In all panels p value ranges are indicated as * $p \leq 0.05$, ** $p \leq 0.01$, *** $p \leq 0.001$ and **** $p \leq 0.0001$.

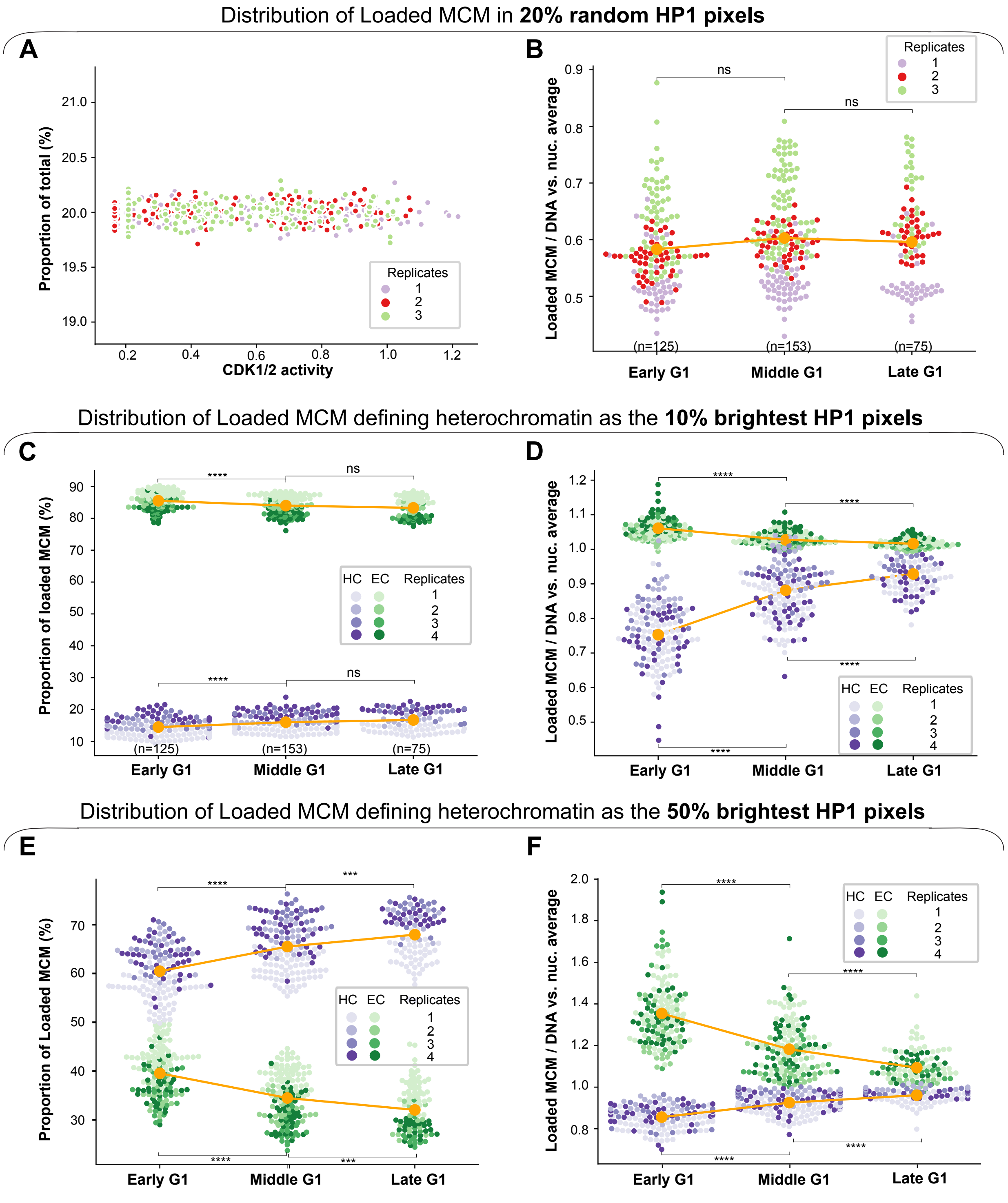


**Supplementary Figure 4. Controls for defining heterochromatin in immunofluorescence images.**

(**A**) Loaded MCM3 in a random 20% selection of pixels relative to CDK1/2 activity in G1 cells plotted as the proportion of total MCM signal in nuclei.

(**B**) Loaded MCM3 concentration in the random 20% selection of pixels in **A** relative to average nuclear loaded MCM3 concentration.

(**C-F**) Quantification of the proportion of loaded MCM3 (**C** and **E**) or the concentration relative to the nuclear average (**D** and **F**) for heterochromatin defined as the 10% or 50% brightest HP1 signals in the indicated G1 subphases. Mean is plotted in orange; one-way ANOVA, Tukey post-hoc test, n (number of cells) is indicated in the figure.

In all panels p value ranges are indicated as * p ≤ 0.05, ** p ≤ 0.01, *** p ≤ 0.001 and **** p ≤ 0.0001.

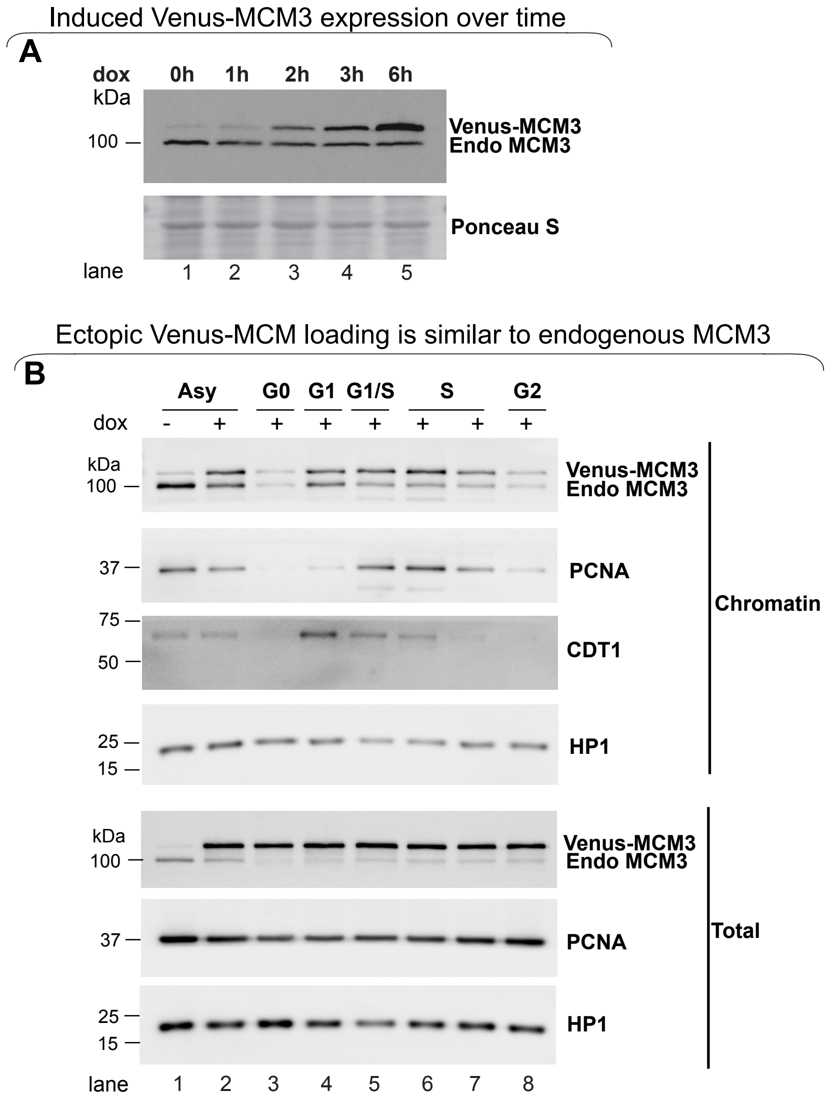


**Supplementary Figure 5. Venus-MCM3 is loaded onto chromatin similarly to endogenous MCM3.**
(**A**) Immunoblot of total protein lysates of asynchronously proliferating RPE1-hTert cells with stably-integrated inducible mVenus-MCM3 treated with 500 ng/ml doxycycline (dox) for 0, 1, 2, 3, or 6 hours.
(**B**) Asynchronous cells or cells synchronized by contact-inhibition (G0) and release were pre-treated with 500 ng/ml dox for 24 hours. Cells were harvested after release at 16, 18, 21,24 and 30 hours to enrich for G1, late G1/early S, mid-S phase, late S and G2 phase respectively respectively. Whole-cell lysates and chromatin fractions were probed for the indicated proteins.

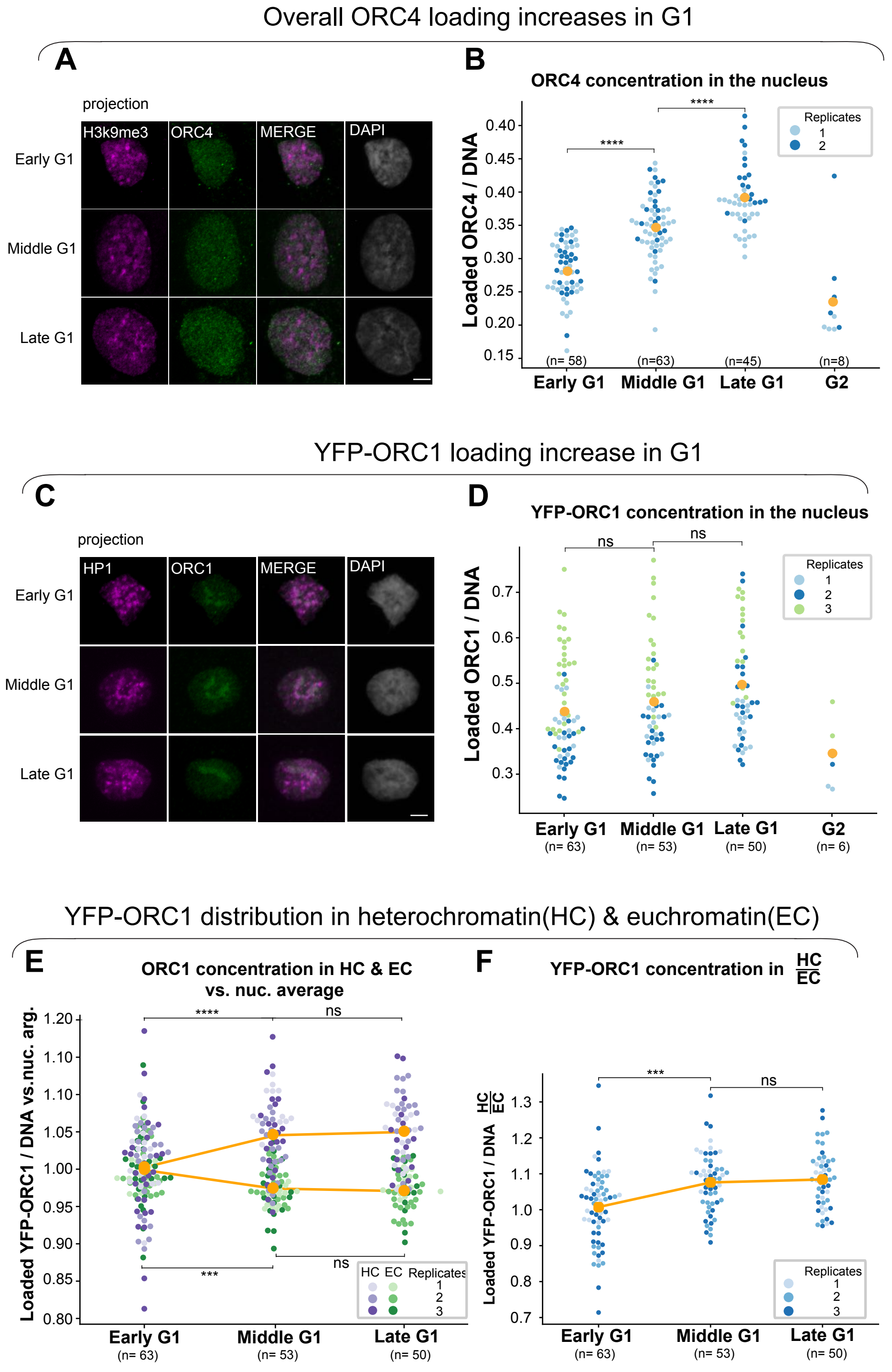


**Supplementary Figure 6. The concentration of loaded ORC increases during G1 phase.**
(**A**) Projections of 3D immunofluorescence images of representative cells after live cell imaging as in Figure 1. Loaded endogenous ORC4 (green) and HP1 (magenta) were detected by immunostaining; scale bar represents 5 μm.
(**B**) Loaded ORC4 concentration normalized to DNA in G1 subphases; two biological replicates are shown; means are plotted in orange. One-way ANOVA, Tukey post-hoc test, n (number of cells) is indicated in the figure.
(**C**) As in **A** except that cells were transfected with plasmid to express dox-inducible YFP-ORC1, treated with 100 ng/ml dox for 48 hours, then extracted, fixed, and stained with GFP antibody; HP1 (magenta), YFP-ORC1 (green), scale bar 5 μm.
(**D**) Quantification of YFP-ORC1 concentration in subphases of G1; two biological replicates are shown; means are plotted in orange.One-way ANOVA, Tukey post-hoc test, n (number of cells) is indicated in the figure.
(**E**) Loaded YFP-ORC1 concentration (YFP signal per unit DAPI) in heterochromatin (purple) and euchromatin (green) relative to the nuclear to average loaded YFP-ORC1 signal; two replicates are shown. One-way ANOVA, Tukey post-hoc test, n (number of cells) is indicated in the figure.
(**F**) Ratio of ORC4 concentration in heterochromatin to euchromatin; two replicates are shown. One-way ANOVA, Tukey post-hoc test, n (number of cells) is indicated in the figure.
In all panels p value ranges are indicated as * p ≤ 0.05, ** p ≤ 0.01, *** p ≤ 0.001 and **** p ≤ 0.0001.

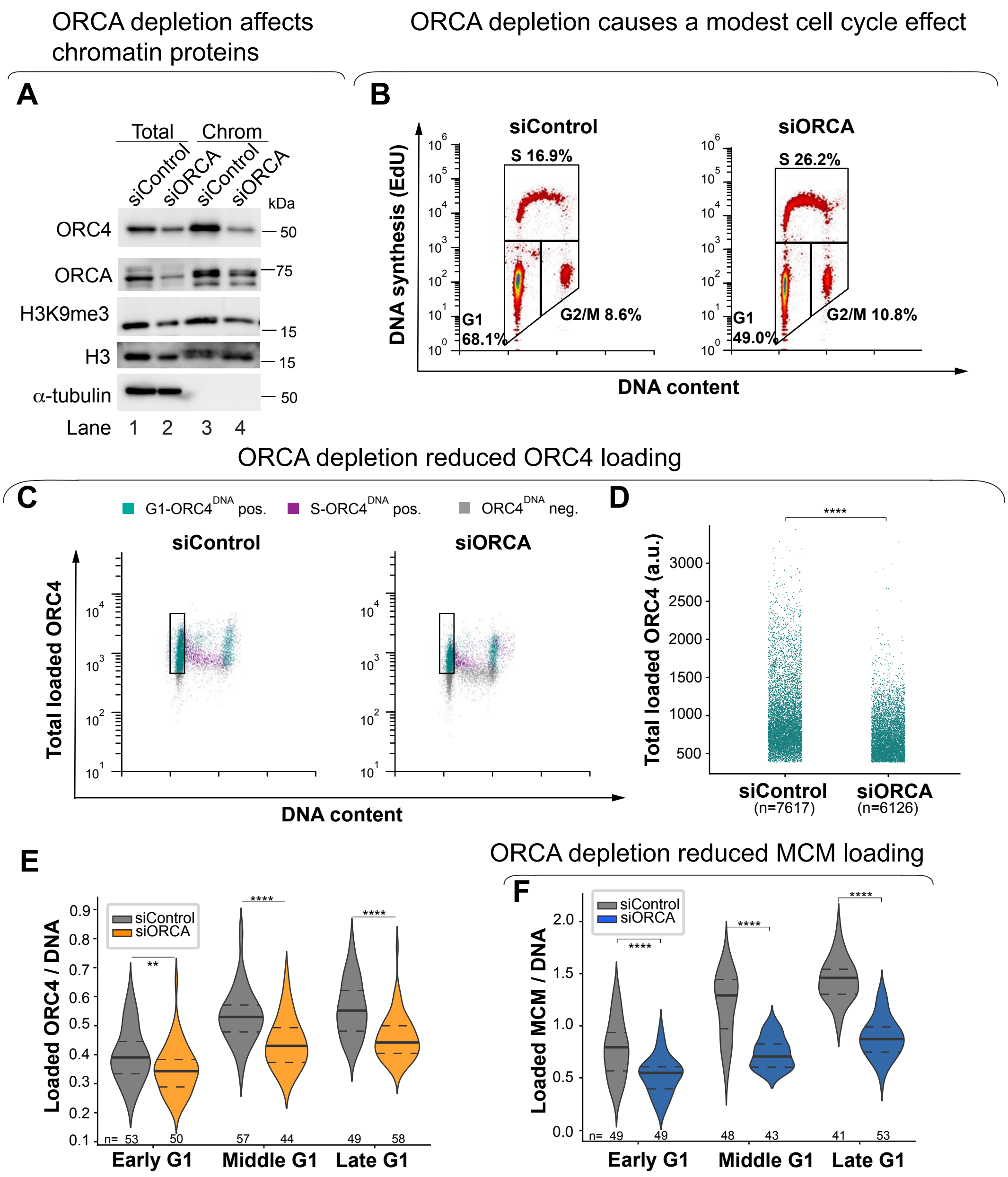


**Supplementary Figure 7. ORCA depletion impairs ORC4 loading and MCM loading.**
(**A**) Asynchronously proliferating RPE1-HTert cells were transfected with 100nM siRNA targeting ORCA or control siRNA for 72 hours before harvesting. Samples were fractionated into DNA-loaded chromatin fraction and total lysate for immunoblotting for the indicated proteins.
(**B, C, D**) Flow cytometry of chromatin-bound proteins in EdU-labeled cells treated as **A**.The distribution of cells in cell cycle phases (**B**) and loaded ORC4 (**C**) is shown. Quantification of loaded ORC4 in the G1 gate (rectangles in C) is shown in (**D**).
(**E, F**) Cells were treated with control siRNA or siRNA targeting ORCA for 48 hours. Loaded ORC4 (**E**) and MCM3 (**F**) immunostaining signals were normalized to DAPI. One-way ANOVA, Tukey post-hoc test, n (number of cells) is indicated in the figure.
In all panels p value ranges are indicated as * $p \leq 0.05$, ** $p \leq 0.01$, *** $p \leq 0.001$ and **** $p \leq 0.0001$.

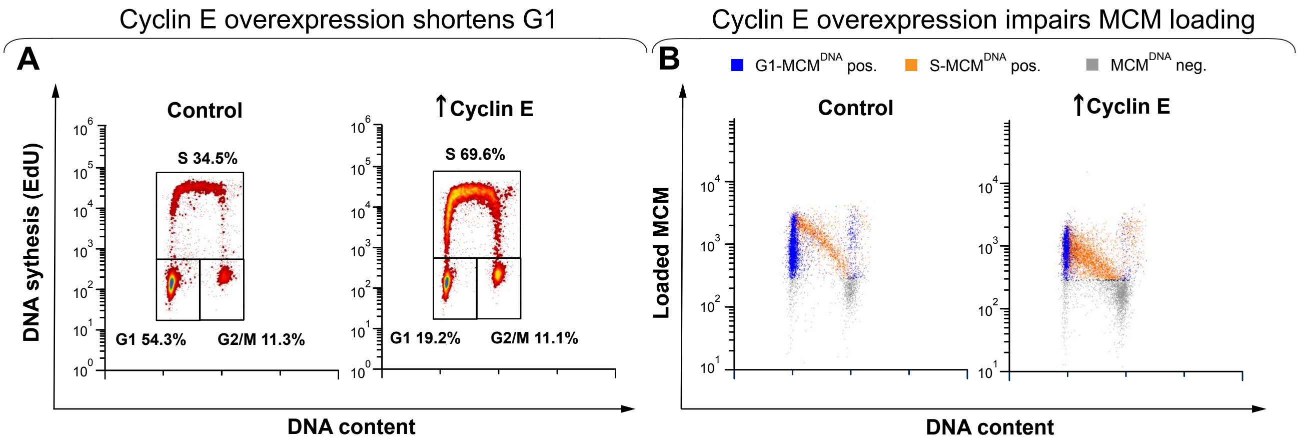


**Supplementary Figure 8. Cyclin E overexpression shortens G1 and induces under-licensing.**
(**A, B**) Flow cytometry of chromatin-bound proteins in EdU-labelled cells treated with 100ng/ml dox to induce ectopic cyclin E expression for 24 hours. The distribution of cell cycle phases (**A**) and loaded MCM (**B**) are shown. Replicate results shown in Matson et al. 2017.psum

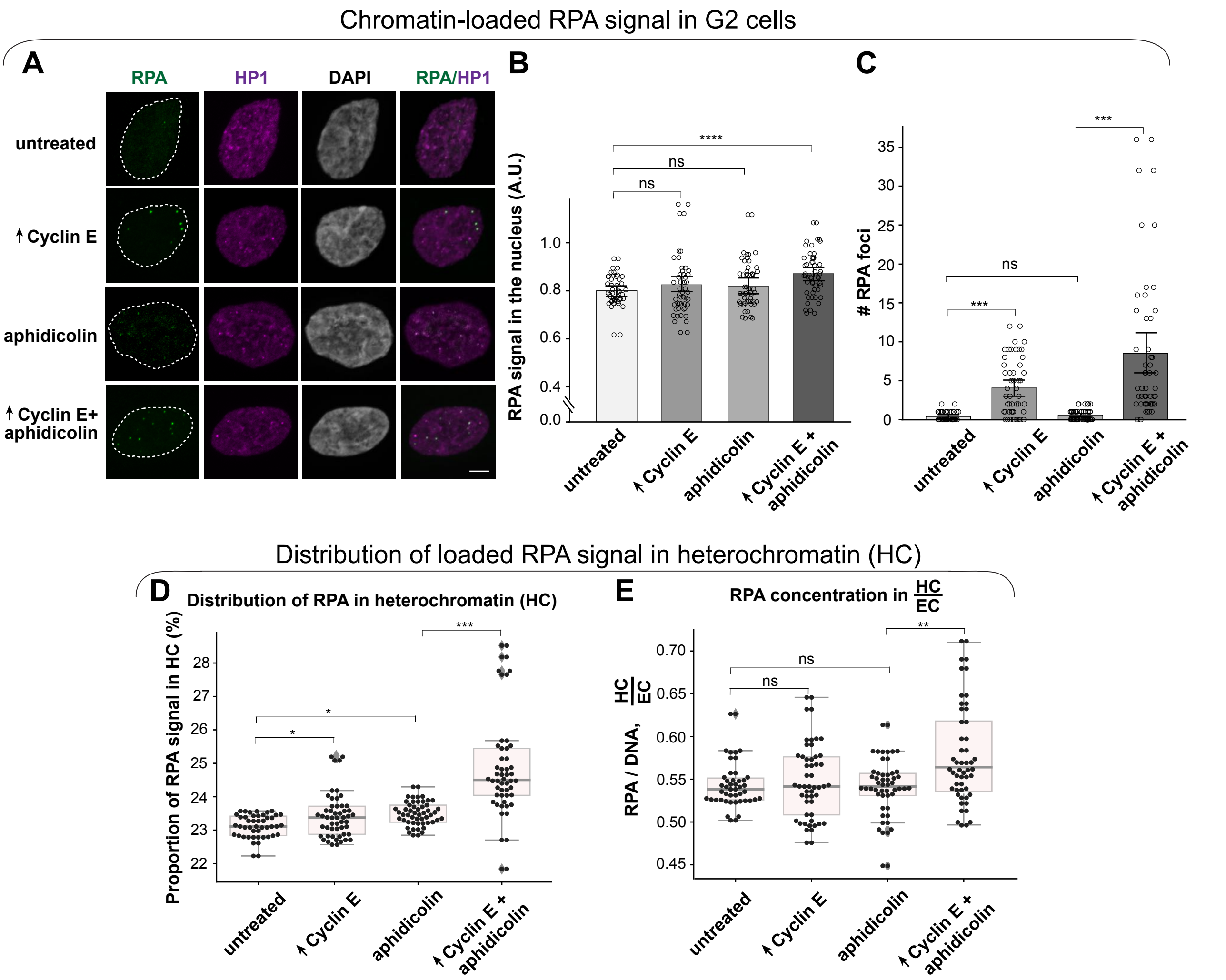


**Supplementary Figure 9. Heterochromatin is more vulnerable than euchromatin to under-replication and DNA damage (similar to 53BP1 analysis in Figure 7)**

(**A**) Projections of 3D immunofluorescence images of representative late G2 cells after live cell imaging as in Figure 1. Cells were treated with 500 ng/ml dox for 18 hours before imaging and with vehicle or 25 ng/ml aphidicolin for the final 4 hours of imaging as indicated. Soluble proteins were extracted prior to fixation and immunostaining for endogenous RPA (green), HP1 (magenta), and DNA (DAPI, grey) scale bar 5 μm. Cells with G1 phases shorter than 4 hours were selected for analysis in the next G2 phase using PCNA variance as an indicator of the S/G2 transition.

(**B, C**) Quantification of total loaded RPA signal in G2 phase nuclei (**B**) and the number of RPA foci per cell (**C**).

(**D**) Proportion of RPA in heterochromatin.

(**E**) Ratio of RPA concentrations in heterochromatin to euchromatin; n (number of cells) >50 for each group; two replicates are shown. In all panels p value ranges are indicated as * $p \leq 0.05$, ** $p \leq 0.01$, *** $p \leq 0.001$ and **** $p \leq 0.0001$.
